# Predestined neutrophil heterogeneity in homeostasis varies in transcriptional and phenotypic response to *Candida*

**DOI:** 10.1101/2022.11.01.514676

**Authors:** Allison K. Scherer, Alex Hopke, Shuying Xu, Adam Viens, Natalie J. Alexander, Kyle D. Timmer, Dakota Archambault, Daniel Floyd, Natalie J. Atallah, Catherine Rhee, Murat Cetinbas, David T. Scadden, Daniel Irimia, David B. Sykes, Ruslan I. Sadreyev, Michael K. Mansour

## Abstract

Once perceived to be homogenous effector cells, neutrophils have since been shown to exhibit population heterogeneity. Here, we established an experimental model of clonal neutrophil heterogeneity using conditionally immortalized clonal granulocyte monocyte progenitors (GMPs) and their mature neutrophil progeny. Transcriptional and epigenetic profiling showed conserved genome-wide signatures of transcription and chromatin accessibility that were specific to individual GMP clones and their paired neutrophil progeny, suggesting that clone specificity is established as early as the GMP stage. Clone-specific genes in vital regulatory pathways were pre-programmed and exhibited delayed expression in the mature neutrophil stage. The clone-specific gene expression in the mature neutrophils paired to enhancer activation in their parental GMPs. To determine whether transcriptional heterogeneity predicted the response to fungal pathogens, neutrophil clones were functionally profiled. Clones demonstrated heterogeneous responses to fungal pathogens *in vitro* and revealed neutrophil subsets with evidence for tailored functional responses to *Candida spp*. as well as specific transcriptional and epigenetic patterns that may explain these differences. Together, this work establishes that heterogenous GMP and neutrophil compartments exist under homeostatic conditions and that these represent predefined clusters that are uniquely adapted to control invasive fungal pathogens.

**Short Summary:** Clonal neutrophil progenitors demonstrate heterogeneity in transcription and chromatin accessibility which may inform response to later fungal challenges.

## INTRODUCTION

Unlike heterogeneity within the adaptive immune cell compartment, evidence for heterogeneity in innate cells has been challenging to define. Within the innate immune compartment, heterogeneous monocyte populations have been identified via cell surface markers and functional profiling (Guilliams et al., 2018; Wolf et al., 2019) and in monocytes and dendritic cells through transcriptional studies (Villani et al., 2017). In neutrophils, the definition of heterogeneity has been difficult despite a variety of approaches to characterization (Qi et al., 2021; Hellebrekers et al., 2017; Filep and Ariel, 2020; Grieshaber-Bouyer and Nigrovic, 2019; Bashant et al., 2021; Deerhake et al., 2021; Mistry et al., 2019; Xie et al., 2020). Transcriptional analysis of murine neutrophils has postulated eight stages of neutrophil differentiation found in the bone marrow and circulation (Xie et al., 2020), while heterogeneity in mature pulmonary neutrophils has been described during *Cryptococcus neoformans* infection (Deerhake et al., 2021). To examine possible heterogeneity in mature neutrophils in response to pathogen challenges, we utilized a well-characterized strategy to isolate and expand clonal neutrophil progenitors, a strategy that permits molecular and functional profiling.

Granulocyte monocyte progenitors (GMPs) have been transcriptionally defined as myeloid committed cells responsible for the production of neutrophils as well as eosinophils, basophils, and monocytes (Kwok et al., 2020). Using a conditionally immortalized GMP model system, we examine clonal populations of neutrophil committed GMPs and their subsequent mature neutrophil progeny (Wang et al., 2006; Sykes and Kamps, 2001). Bone marrow GMPs were harvested, conditionally immortalized using enforced expression of Hoxb8, and single cells sorted to generate clonal populations. Sequential testing of these GMPs identified consistently neutrophil-fated GMP clones which could be assayed for heterogeneity through functional, transcriptional, and epigenetic profiling.

Neutrophils are the most abundant circulating immune cell and have a short lifespan ranging from a few hours to a few days (Tak et al., 2013; Basu et al., 2002; Pillay et al., 2010). During this time, they perform a variety of roles in the setting of injury and disease, and the loss of neutrophils is a significant risk factor for developing an invasive fungal infection. Here, we examined clonal neutrophils for evidence of functional heterogeneity in response to *Candida spp*. including *Candida albicans, Candida glabrata*, and *Candida auris* which account for most occurrences of invasive candidiasis in the United States (Ricotta et al., 2020). Following neutrophil generation from the clonal GMP cell lines, we systematically tested clone-to-clone differences *in vitro* by observing the neutrophil response to *Candida spp*. Clones maintained a consistent functional phenotype and demonstrated distinct neutrophil responses to fungal pathogens.

To address potential subpopulation heterogeneity between the GMP and neutrophil states, we characterized multiple clones as immature GMPs as well as terminally differentiated neutrophils via transcriptional, epigenetic, and functional profiling. Transcriptional and epigenetic analysis of paired clones highlights conserved and closely associated gene expression between the GMP stage and the mature neutrophil stage. The clones followed the same maturation trajectory while maintaining key differences in their transcriptome. Functionally, clones clustered into distinct groups in response to *Candida*. Taken together, our data suggest the existence of heterogeneous populations of neutrophils with variable responses to the human fungal pathogen, *Candida*. Future characterization will lead to an understanding of the molecular mechanisms that neutrophils require to best combat fungal pathogens and may point towards future therapeutic interventions to tailor a patient’s innate immune response in the setting of infection.

## RESULTS

### Conditional immortalization of granulocyte monocyte progenitors for clonally derived neutrophils

Heterogeneity of the granulocyte monocyte progenitors (GMPs) and their ensuing neutrophils have been defined in several ways (Scherer et al., 2021). **Figure 1a** illustrates potential mechanisms through which neutrophil heterogeneity may arise: 1) GMPs may be homogeneous in the bone marrow compartment and result in mature neutrophils which respond equally to stimuli, 2) neutrophils arise from heterogeneous GMPs and maintain distinct characteristics through maturation, or 3) GMPs are homogenous but result in heterogeneous neutrophils which respond differently to various stimuli. To test these hypotheses, we leveraged a tool to generate clonally derived neutrophils with which to examine baseline transcriptional and epigenetic characteristics, as well as variation in effector functions to *Candida* challenge (**Figure 1b**).

**Figure 1.**
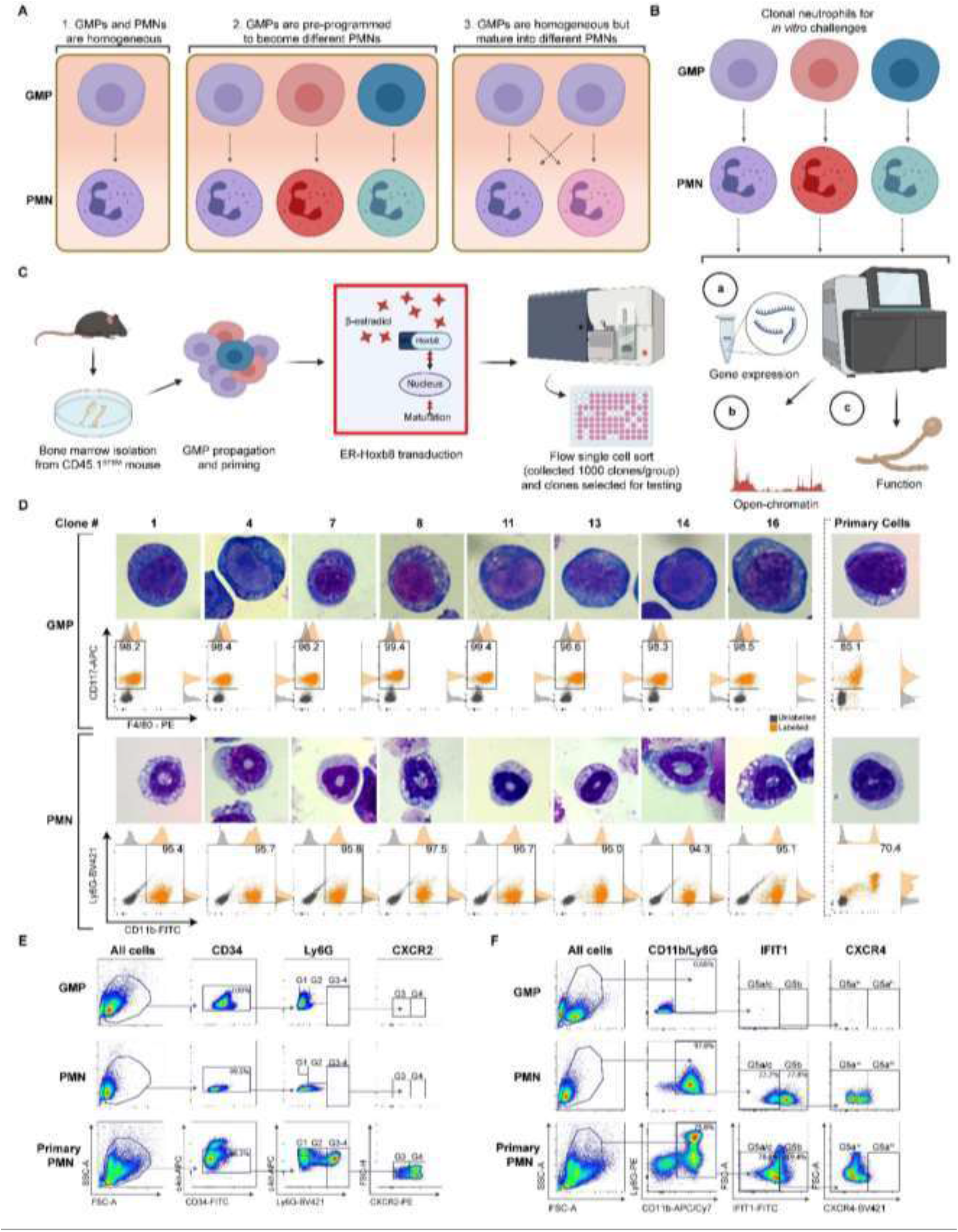
Generation and maturation characterization of clones. **(a)** Illustration of potential areas of neutrophil heterogeneity where 1) homogeneous populations of granulocyte monocyte progenitors (GMPs) mature into homogeneous neutrophils (PMNs) which respond equally to stimuli, 2) GMPs are pre-programmed to mature into neutrophils which respond to specific stimuli, or 3) GMPs belong to a homogeneous population which become heterogeneous during neutrophil maturation (adapted from (Scherer et al., 2021) with BioRender.com. **(b)** Outline of the experimental method in which clonal GMPs were kept in culture, matured in the absence of β-estradiol, and then subsequentially used for transcriptional and epigenetic studies or challenged with a fungal pathogen. **(c)** Schematic which outlines construction of GMP clones and downstream clonally derived neutrophils (panels a-c created with BioRender.com). **(d)** Representative Wright Giemsa staining and cell surface labelling for GMP to PMN clones used in later RNA-seq and ATAC-seq analysis. For each clone, representative flow plots are given demonstrating the maturation state where grey dots indicate the unstained control sample, and the orange dots indicate the stained sample for each GMP and PMN. Plots to the right indicate cells harvested from murine bone marrow for comparison. **(e)** Further characterization by cell surface markers from a representative clonal GMP to immature neutrophil panel (G0-4), and **(f)** its corresponding mature neutrophil labelled as G5a, G5b, or G5c as previously described (Xie et al., 2020). Cells harvested from murine bone marrow are shown for comparison.

Previous work to define neutrophil heterogeneity has examined primary GMP and neutrophil single cells with single-cell transcriptomics (Deerhake et al., 2021; Xie et al., 2020). These data support neutrophil heterogeneity in health and disease states but are limited in their ability to examine neutrophil effector functions or epigenetic markers in additional experiments. To circumvent this obstacle, we conditionally immortalized GMP cells from CD45.1^STEM^ mice (Mercier et al., 2016) using a truncated estrogen receptor homeobox b8 (ER-Hoxb8) construct (Wang et al., 2006; Sykes and Kamps, 2001). Bone marrow GMPs were harvested and transduced with MSCV-based retrovirus encoding ER-Hoxb8 and single-cell clones were expanded in 96-well plates (**Figure 1c**). Of those, 16 robust neutrophil-fated GMPs were selected for characterization.

Selected clones matured into neutrophils following the removal of β-estradiol from culture media, as demonstrated by Wright Giemsa staining and cell surface marker expression (**Figure 1d**). In the presence of β-estradiol, all clones were similar, appearing as GMPs as characterized by large mononuclear cells in Wright Giemsa staining and by high CD117+ expression in comparison to their paired matured neutrophil clone. Neutrophils derived after the removal of β-estradiol from the media were confirmed by CD11b+ expression. Although the neutrophils show weak Ly6G labeling in comparison to primary murine neutrophils harvested from bone marrow, the clones demonstrate the expected neutrophil morphology by Wright Giemsa staining. This low protein expression of Ly6G has been noted previously (Negoro et al., 2020) and appears to be an artifact of *in vitro* culture; Ly6G expression can be boosted with stimulation by GM-CSF and IL-4 (Fites et al., 2018).

To demonstrate that the GMP clones mature similarly to primary neutrophils, we utilized previously described flow panels for known cell surface markers (Xie et al., 2020). Here, primary mouse samples from the bone marrow indicate maturation stages (Xie et al., 2020) including G1 (promyelocytes, CD117^+^Ly6G^−^), G2 (metamyelocytes, CD117^+^Ly6G^+/−^), G3 (immature neutrophils, CD117^+/−^Ly6G^+^ CXCR2^−^), or G4 (immature neutrophils, CD117^+/−^Ly6G^+^CXCR2^+^) (Xie et al., 2020). Our clonal GMPs expressed markers consistent with the G0 (GMP) phase (**Figure 1e, Suppl. Fig. 1–2)**, and neutrophils differentiated *in vitro* lost these early maturation cell markers (**Figure 1e**). We also examined the expression of markers associated with G5a-c (G5a: IFIT1^−^CXCR4^lo^, G5b: IFIT1^+^, G5c: IFIT1^−^CXCR4^hi^) mature type neutrophils (Xie et al., 2020). Here, our neutrophil clones closely resemble G5b cells (77.2%) with IFIT1 expression and G5a cells (22.2%) with some CXCR4 expression (**Figure 1f, Suppl. Fig. 3**). Some overlap in the G5a and G5b populations is expected as these stages are phenotypically quite similar (Xie et al., 2020).

### Transcriptional profiling reveals GMP heterogeneity that predicts neutrophil heterogeneity

To understand the baseline transcriptional landscape, we selected a subset of unstimulated, matched GMP clones and their neutrophil progeny. Clones showed an overall similar transcriptional profile within the GMP and neutrophil states, though individual clones had several differentially expressed genes (DEGs) between each other at both the progenitor and mature cell stages (**Figure 2a)**. Interestingly, although the strongest separation was observed between progenitors and mature cells (dimension 1 in **Figure 2b**) there was also a marked distinction among expression profiles of individual clones regardless of the stage (dimension 2 in **Figure 2b**). Here, the replicates appear closely related to each other (*e.g*., blue GMP open circles or blue neutrophil closed circles) and the GMP state shows expression similarity with the clone’s paired neutrophil state (*e.g*., the dotted lines between blue GMP open circles in relation to blue neutrophil closed circles). These data highlight that the clone-specific expression pattern of GMPs is associated with neutrophil gene expression and potentially indicates that neutrophil heterogeneity already exists within the GMPs.

**Figure 2.**
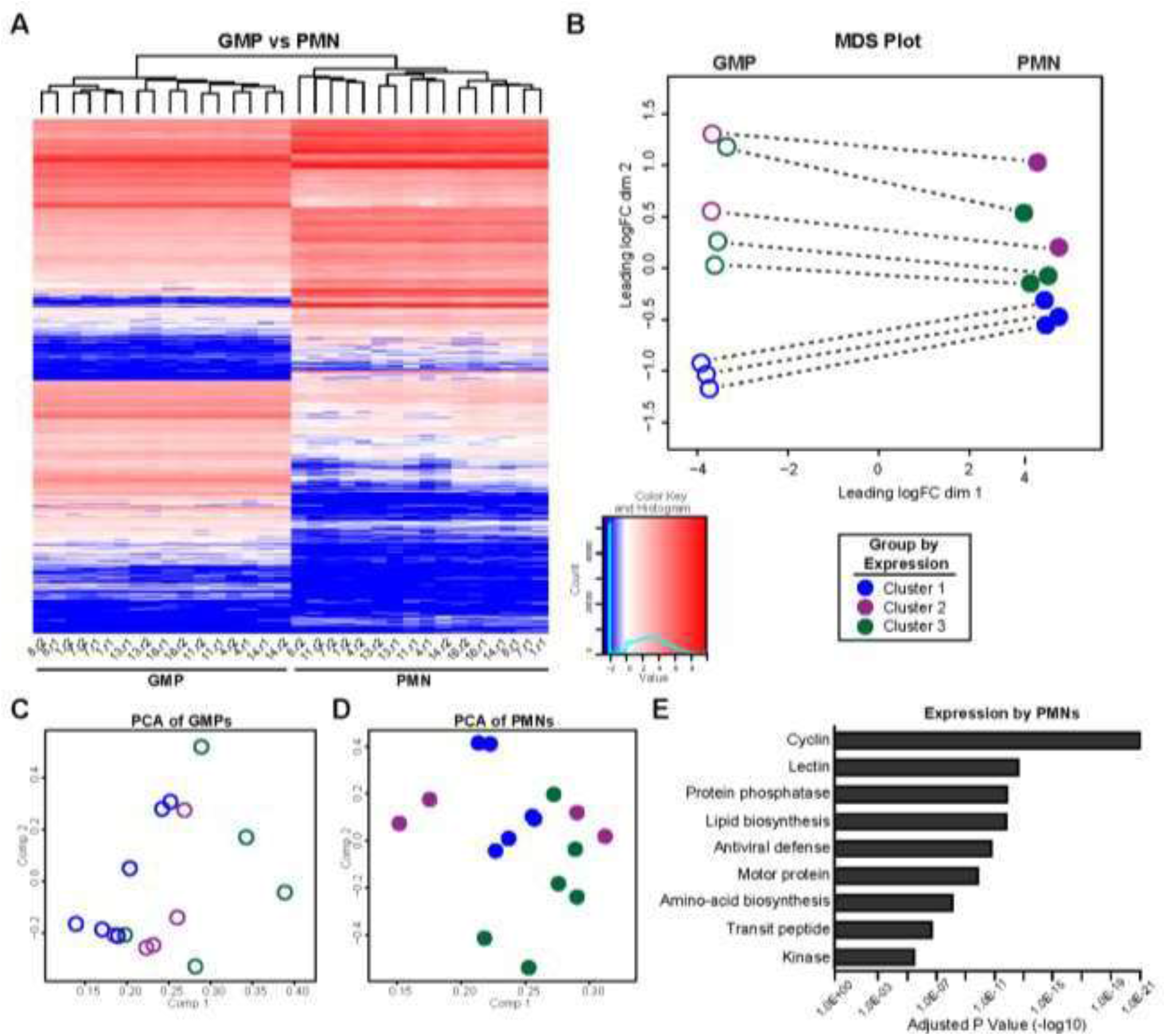
Neutrophil heterogeneity pattern identified through transcriptional profiling. Naïve paired GMP and neutrophil clones were used to identify the clone’s baseline transcriptional profile. **(a)** Heatmap of expression values (RPKM in log2 scale) for the differentially expressed genes (DEGs) in all pairs of respective GMP and neutrophil clones. **(b)** MDS plot of the top 500 most varying genes between GMPs and neutrophils. The dotted line indicates the GMP (open circles) to neutrophil (closed circles) transition for each clone and colors indicate the clone’s transcriptional group. **(c, d)** PCA plots demonstrating the grouping of GMPs **(c)** and neutrophils **(d)** based on gene expression. **(e)** Functionally enriched pathways identified by the DAVID tool among DEGs between neutrophil clones. (granulocyte monocyte progenitor, GMP; neutrophil, PMN)

To illustrate potential relationships between clones, we clustered the eight clones based on their transcriptional profiles. Internal clustering of the clones based on their gene expression is shown in the GMPs (**Figure 2c**) and neutrophils (**Figure 2d**). Clones separated into three expression-based clusters: i) 1, 7, 8 (blue), ii) 11, 13 (purple), and iii) 4, 14, 16 (green) at the GMP level (**Suppl. Fig. 4a**), with a total of 860 inter-cluster DEGs, and at the neutrophil level (**Suppl. Fig. 4b**), with a total of 523 inter-cluster DEGs. Notable similarities between clones using transcriptional clustering are seen in clones 7 and 8 (blue), 11 and 13 (purple), and 14 and 16 (green). These clones appear to be more closely related at the mature neutrophil state than at the GMP state (**Figure 2d)**, suggesting the clustering based on gene expression can partially be used to define neutrophil subpopulations.

Using the Database for Annotation, Visualization, and Integrated Discovery (DAVID) (Huang et al., 2009), we identified enriched pathways of interest among these DEGs for both GMPs and neutrophils (**Figure 2e**). We compared the GMP clones to each other and the neutrophil clones to each other to obtain DEGs. We identified the union of DEGs for the expression by GMPs and neutrophils, separately. Unsurprisingly, pathways related to immunity and innate immunity were included in both GMP and neutrophil gene expression. Gene expression by neutrophils showed several pathways relevant to fungal infection (**Figure 2e**). For example, lectins were identified which are critical cell surface markers used by neutrophils to identify and eliminate fungal pathogens (Gazendam et al., 2014; Urban and Backman, 2020). Kinase pathways were also enriched which may contribute to later neutrophil effector functions (*e.g*., spleen tyrosine kinase (Negoro et al., 2020)). Overall, the clone’s transcriptional signatures suggest that heterogeneity at the GMP level can translate into later heterogeneity in the mature neutrophil. This points towards the second hypothesis proposed in Figure 1a in which GMPs are pre-programmed, thereby resulting in mature neutrophil heterogeneity.

### Epigenetic signatures predict transcriptional expression clusters

To assess the chromatin state for clone-specific epigenetic differences, we performed ATAC-sequencing (Buenrostro et al., 2015) on the paired GMP and neutrophil clones. **Figures 3a-b** show examples of genomic tracks for genes with differences in chromatin accessibility between GMPs (**Figure 3a**) and neutrophils (**Figure 3b**). Several interesting genes for neutrophils arose which play important roles in response to pathogens, including Clec7a, Alox5, and CD177. Of note, Clec7a encodes for Dectin-1 which recognizes β-glucans in the *Candida spp*. cell wall (Gazendam et al., 2014) and is Syk-dependent. Similarly, Alox5 is responsible for the translocation of 5-lipoxygenase to the nuclear envelope membrane where it converts arachidonic acid into leukotriene A4 and later leukotriene B4 (Poplimont et al., 2020), and CD177 is specific to granules and mobilized to the cell surface with cell activation (Grieshaber-Bouyer and Nigrovic, 2019). Variability in open chromatin regions presents another potential method for identifying potential neutrophil subpopulations.

**Figure 3.**
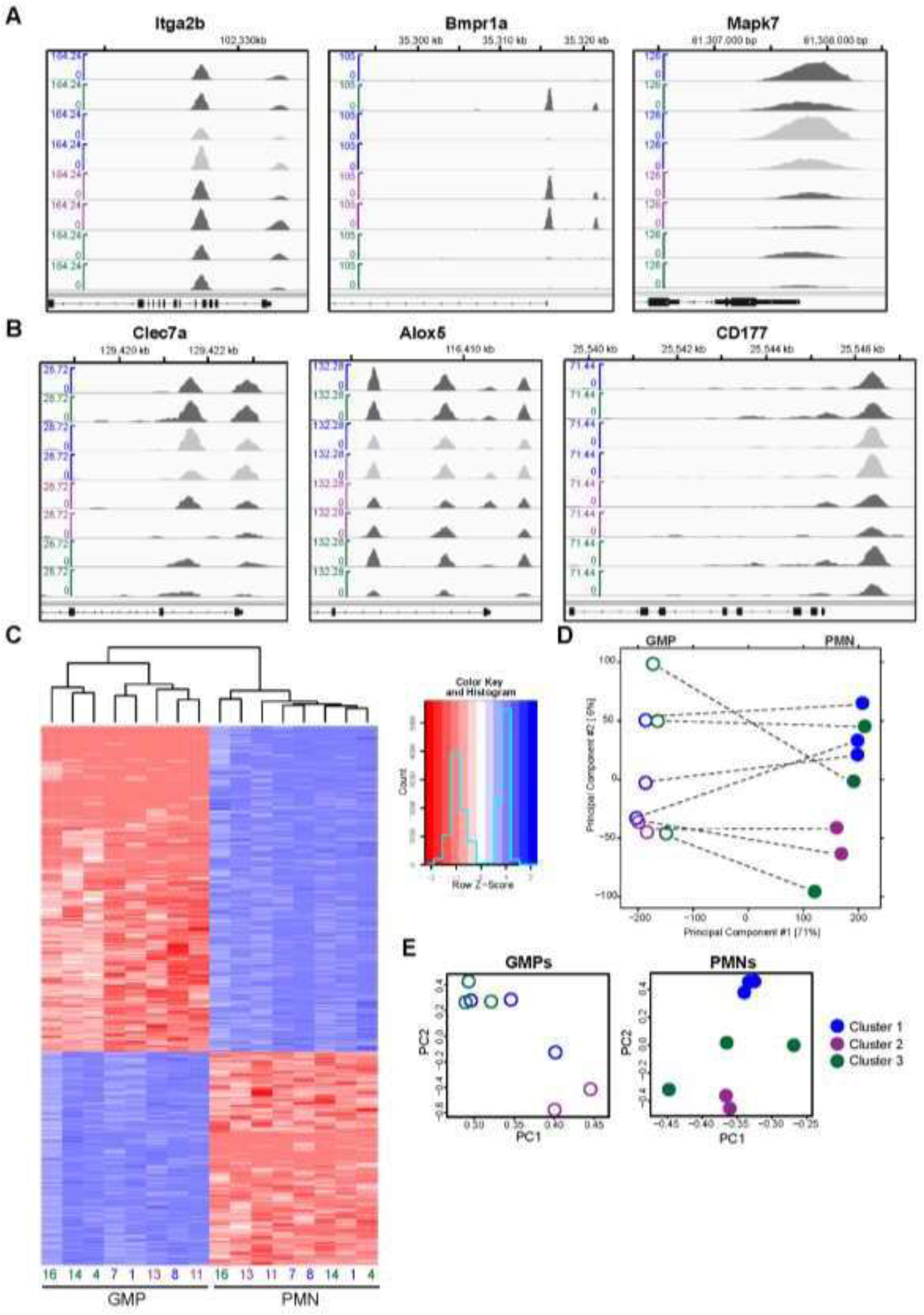
Genome-wide patterns of chromatin accessibility demonstrate similarities of neutrophil clone grouping in transcriptional clusters. **(a, b)** Examples of genomic ATAC-seq tracks in the vicinity of selected promoters and enhancers for GMPs **(a)** and neutrophils **(b)**. **(c)** Heatmap of chromatin accessibility for each of the differentially accessible regions (DARs) between GMPs and neutrophils. Chromatin accessibility at each DAR is shown as Z-scores relative to the distribution of ATAC-Seq signals across all clones for the given DAR. **(d)** PCA plot of paired GMP and neutrophil clones based on the union of DARs between each pair of GMPs and neutrophils. The dotted line indicates the GMP (open circles) to neutrophil (closed circles) transition for each clone; colors indicate the clone’s previously defined transcriptional cluster. **(e, f)** Separate PCA plots for GMP **(e)** and neutrophil clones **(f)** based on DARs among clones of the same cell type indicating a correlation between the clone’s previously defined transcriptional color-coded clusters. (granulocyte monocyte progenitor, GMP; neutrophil, PMN)

A comparison of the GMP to the neutrophil clones revealed 29,966 differentially accessible regions (DARs) between GMPs and neutrophils, from a total of 74,009 consensus peaks of chromatin accessibility. The heatmap in **Figure 3c** illustrates chromatin accessibility at each of these DARs across clones. Differences in the epigenetic profiles between GMPs and neutrophils are the strongest, but there are also smaller differences in the epigenetic signatures between individual clones at each cell stage (**Figure 3c**). The PCA plot illustrates the similarities of chromatin accessibility patterns across GMP and neutrophil clones (**Figure 3d**). Most clones appear to have some correlation between the GMP and the neutrophil stage (*e.g*., open purple circles to closed purple circles), as was found with transcriptional profiling (**Figure 2b**), however, other clones appear to be less correlated (*e.g*., open green circles to closed green circles).

Promoter chromatin accessibility showed a distinct yet modest correlation with gene expression in the same clone (**Figure 3**, **Suppl. Fig. 5a-b**). However, the comparison of promoter accessibility and gene activity for the same gene across multiple clones showed a strong correlation. **Suppl. Fig. 5c** shows the distribution of Pearson correlation coefficients between gene expression and promoter chromatin accessibility across all clones calculated for each individual gene across the genome, whereas **Suppl. Fig. 5d** shows the same distribution for all intergenic ATAC-seq peaks as potential enhancers and the expression of the closest nearby gene in a 1 Mb vicinity. A large fraction of promoters and enhancers show a strong quantitative correlation with the expression level of the corresponding gene across all individual clones. Together, these epigenetic signatures suggest that GMPs have enhancer activation that precedes neutrophil maturation which may alter the transcriptional and phenotypic response of the mature neutrophil.

### Clones follow the same differentiation trajectory while maintaining specific expression differences

We sought to understand the extent to which the signatures of expression differ between individual clones at the GMP stage translated into expression differences at the neutrophil stage. The heatmap in **Figure 4a** shows the extent of overlap between clone-specific DEGs observed for individual clone pairs at the GMP stage and clone-specific DEGs observed for the same clone pairs at the neutrophil stage. This overlap is shown as a fraction relative to GMP DEGs (upper part of the heatmap) and to neutrophil DEGs (lower part of the heatmap). Although this overlap was relatively modest (typically 10-30%), it was highly statistically significant (**Suppl. Table 1)**, especially in the context of strong transcriptional changes that accompany differentiation. Remarkably, all clones shared a strong similarity of their differentiation gene signatures (**Figure 4b, Suppl. Table 2**). Approximately ~80% of genes differentially expressed between GMPs and neutrophils were shared between all clones (**Figure 4b**). When we analyzed the subset of these shared DEGs that have a DAR in either a nearby promoter or enhancer region, we found that a large fraction of these genes showed a strong correlation between the levels of chromatin accessibility and gene expression.

**Figure 4.**
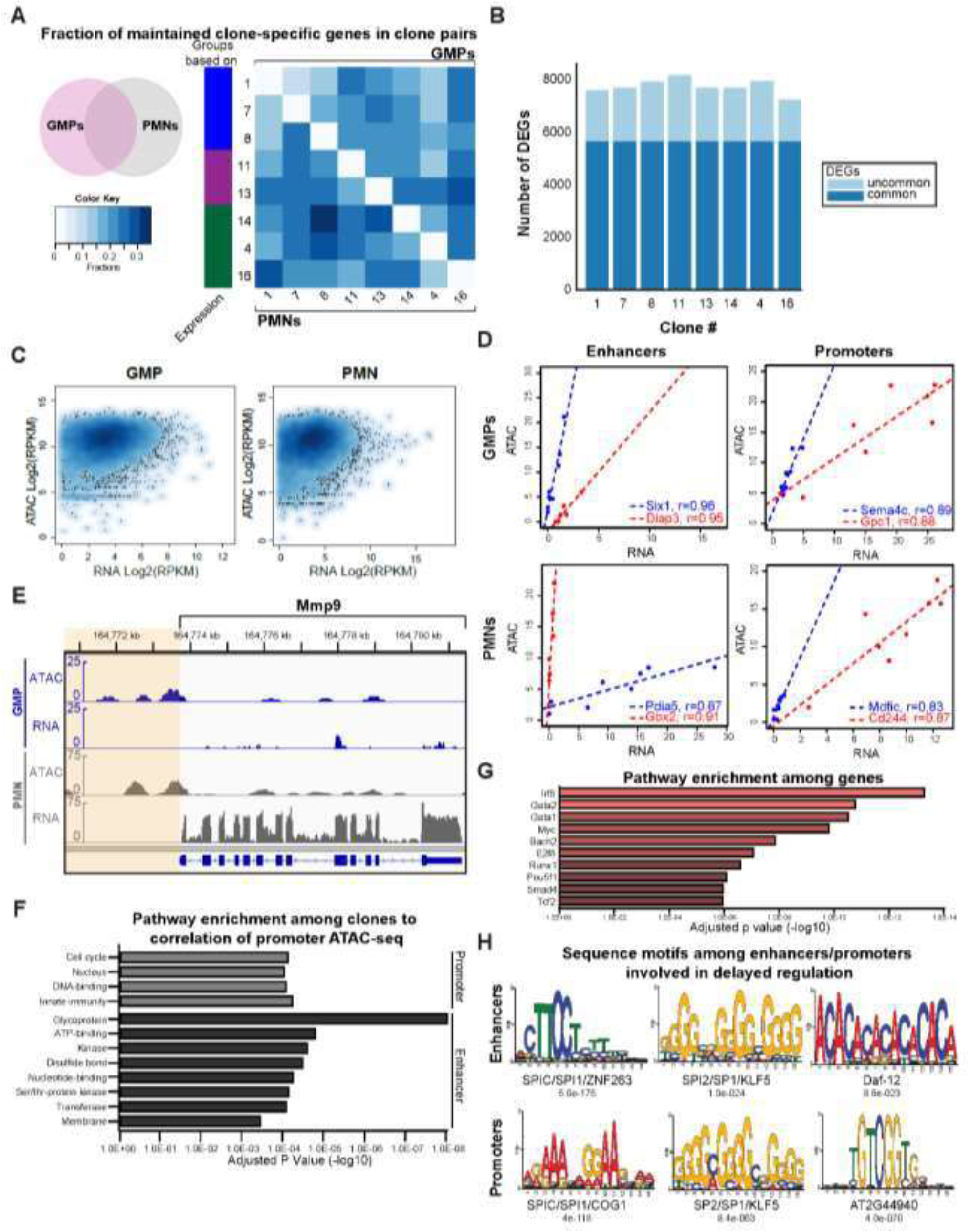
Concerted regulation of clone-specific gene expression and chromatin accessibility during GMP to neutrophil differentiation. **(a)** Signatures of transcriptional differences between individual clones at the GMP stage are partially but significantly maintained at the PMN stage. The set of differentially expressed genes (DEGs) for each pair of GMP clones was overlapped with the set of DEGs for the corresponding pair of PMNs (see a schematic Venn diagram on the left). The heatmap shows the degree of overlap between these clone-specific DEGs at the GMP and PMN stage. For each clone pair, this degree of GMP/PMN overlap is shown as a fraction of overlapping DEGs relative to the total number of DEGs at the GMP stage (above diagonal) and at the neutrophil stage (below diagonal). **(b)** The majority of differentiation-associated transcriptional changes are common among all clones. Barplot of DEG numbers between the GMP and neutrophil stage of each clone, with the number of DEGs shared among all clones shown in dark blue. **(c)** When analyzed across the genome in the same clone, gene expression has a modest correlation with chromatin accessibility at the promoters. Scatter plot of RNA-seq vs promoter ATAC-seq signals (RPKM in log2 scale) for all genes across the genome. **(d)** Chromatin accessibility of a single promoter or enhancer across all clones often shows a strong correlation to the expression of the closest gene. Examples of scatterplots for the correlation between gene expression and chromatin accessibility of a given enhancer (left) or promoter (right) in GMPs (top) and neutrophils (bottom). Each point represents a single clone. Linear regression lines and Pearson R^2^ values are indicated in each plot. **(e)** Example of genomic tracks for enhancers whose clone-specific chromatin accessibility at the GMP stage had a delayed manifestation in clone-specific expression of a nearby gene only at the neutrophil stage of the same clone. **(f)** Genes with the strongest correlation of expression to promoter or enhancer chromatin accessibility are enriched in specific functional pathways. Top enriched pathways according to DAVID pathway enrichment analysis among the top 500 genes whose expression was most correlated to promoter (top) or enhancer (bottom) chromatin accessibility. **(g)** Functional pathways and targets of DNA-binding factors enriched among the genes with a clone-specific regulation delayed with respect to an earlier clone-specific chromatin accessibility of nearby enhancers. Enrichment estimates by the EnrichR tool. **(h)** Analysis of sequence motifs at the corresponding enhancers and promoters suggest the involvement of key transcription factors in this delayed regulation.

**Table 1.** Cell surface marker antibody panels.

| Panel | Antigen | Company | Catalog # | Fluorochrome | RRID | Clone | Reference |
| --- | --- | --- | --- | --- | --- | --- | --- |
| **G1-G4** | **CD117 (c-kit)** | Biolegend | 105812 | APC | AB_313221 | 2B8 | (Xie et al., 2020) |
|  | **CD34** | Invitrogen | 11-0341-85 | FITC | AB_465021 | RAM34 |  |
|  | **Ly6G** | BioLegend | 127628 | BV421 | AB_2562567 | 1A8 |  |
|  | **CXCR2** | BioLegend | 149303 | PE | AB_2565691 | SA044G4 |  |
| **G5a-c** | **CD45.1** | BioLegend | 110714 | APC |  | A20 |  |
|  | **CD11b** | BioLegend | 101226 | APC/Cy7 | AB_830642 | M1/70 |  |
|  | **Ly6G** | BioLegend | 127608 | PE | AB_1186099 | 1A8 |  |
|  | **IFIT1** | Sigma-Aldrich | ABF117 | -- |  | -- |  |
|  | **FITC** | Invitrogen | A32731T R | FITC | AB_2866491 | -- |  |
|  | **CXCR4** | BioLegend | 146511 | BV421 | AB_2562788 | L276F12 |  |
| **GMP** | **Lin-Biotin** | BioLegend | 133307 | -- | AB_11124348 | Ter-119, M1/70, RB6-8C5, 145-2C11, RA3-6B2 | (Challen et al., 2009) |
|  | **Lin-SA** | BioLegend | 405241 | BV711 | AB_11124348 | -- |  |
|  | **CD117** | BioLegend | 105812 | APC | AB_313221 | 2B8 |  |
|  | **Ly6A/E** | BioLegend | 108128 | BV421 | AB_2563064 | D7 |  |
|  | **CD16/32** | BioLegend | 101318 | PE/Cy7 | AB_2104156 | 93 |  |
|  | **CD34** | Invitrogen | 11-0341-85 | FITC | AB_465021 | RAM34 |  |
|  | **CD45.2** | BioLegend | 109824 | APC/Cy7 | AB_830789 | 104 |  |

When we compared the levels of gene expression and chromatin accessibility of the corresponding promoter across all genes between GMP and neutrophil stages of the same clone to the magnitude of the corresponding changes in promoter chromatin accessibility for the same gene, we found only a modest genome-wide correlation (**Figure 4c**). However, there was significant correlation when one focused on the promoter of a single gene or a single enhancer-gene pair and compared chromatin accessibility with gene expression across multiple clones. Illustrated in **Figure 4d** are examples of enhancers and promoters which have a strong correlation between chromatin accessibility and expression of the closest gene across GMPs or PMNs. Together, this suggests that the crosstalk between clone-specific regulation of gene expression and chromatin remodeling may play a role in maintaining clonal integrity across the maturation transition.

Next, we analyzed the temporal progression of clone-specific patterns of chromatin accessibility and gene expression between GMPs and neutrophils. Notably, we found that several genes had a temporal delay between the onset of clone-specific chromatin accessibility of a nearby enhancer and clone-specific expression of the gene. Some genes had clone-specific differences in the ATAC-seq signal at a nearby enhancer at the GMP stage that did not manifest in a significant clone-specific difference in the corresponding gene expression. However, this clone-specific expression difference was manifested later, at the neutrophil stage. Examples of genomic ATAC-seq and RNA-seq tracks for Prg3 (**Figure 4f)** show the temporal development of clonal differences in gene expression at the neutrophil stage following the onset of clonal differences in enhancer chromatin accessibility at the GMP stage. Functional enrichment analyses (Huang et al., 2009) suggested the regulation of central pathways (**Figure 4f**) relevant to innate immunity and kinase activity were noted in the results for promoters and enhancers, respectively. Here, innate immunity pathways include the cell’s ability to recognize pathogens and activate effector functions which suggest that clones already differ in the unstimulated naïve state in their ability to regulate these pathways. Kinases are particularly important for neutrophil recruitment, recognition, and cellular activation of effector functions. As an example, we have previously demonstrated that Syk is critical for neutrophil recognition, phagocytosis, and elimination of *Candida spp*. (Negoro et al., 2020).

EnrichR analysis (Chen et al., 2013; Kuleshov et al., 2016; Xie et al., 2021) pointed to the enrichment of targets of key transcription factors involved in development and hematopoiesis (**Figure 4g**). Of interest, interferon regulatory factor 8 (Irf8), a known regulator of myelopoiesis in mice and humans (Yáñez et al., 2015), was an enriched target. Irf8 acts downstream of lineage commitment to control neutrophil production (Yáñez et al., 2015) and may play a role in defining a sub-group of neutrophils. The transcription factors GATA-binding protein (GATA) −1 and GATA-2 are involved earlier in hematopoiesis. GATA-1 influences hematopoietic stem cells (HSC) to differentiate into megakaryocyte-erythroid progenitors, and GATA-2 in connection with PU.1 influences HSCs to differentiate into granulocyte monocyte specific precursors (Friedman, 2007; Johnson et al., 2020; Rodrigues et al., 2008). Together, these enriched transcription factors indicate a possible connection to delayed regulation in differentiation.

MEME (Bailey et al., 2015) analysis of sequence motifs enriched at the enhancers whose clone-specific chromatin regulation preceded clone-specific gene expression, as well as the sequence motifs at the promoters of these genes, demonstrated the enrichment of transcription factor binding sites (**Figure 4h**). Notably, Spi-1 was found within both enhancers and promoters. Spi-1 codes for PU.1, a transcriptional regulator of hematopoietic stem cell differentiation including neutrophils (Li et al., 2020). Alterations in this transcriptional regulator would be expected to impact GMPs on a clonal level. Additionally, Specificity protein 1 (Sp1) is a well-known transcription factor that actively competes with Sp3 for DNA binding resulting in the enhancing or repressing of gene expression (Vizcaíno et al., 2015). The Sp family are implicated in cell growth, differentiation, and apoptosis (Vizcaíno et al., 2015) suggesting Sp1 could also explain some of the clonal maturation differences.

### Transcriptional and epigenetic signatures inform clone response to fungal challenge

Neutrophils perform a variety of host functions including the recognition of pathogens, recruitment to infection sites, and containment or clearance of invasive organisms. To determine if the clones had differences in baseline transcriptional signatures in relation to these functions, we highlighted genes involved in cell trafficking, signaling pathways, and pattern-recognition such as toll-like receptors (TLRs) (**Figure 5a**). Although most highlighted genes appear similar in transcription, notable genes with alternate profiles include Tlr9 and Tlr1 for pattern recognition and Alox5 and Alox15 for neutrophil chemotaxis and swarming. We further examined spleen tyrosine kinase (syk), a critical kinase for neutrophil effector functions during a fungal challenge, as an example of the relationship between neutrophil gene expression, open chromatin, and function. The clones demonstrate similarities in their transcriptional (**Figure 5b)** and epigenetic (**Figure 5c)** profile of syk and pattern closely to the clusters identified in **Figure 2**. To determine if the combined transcriptional and epigenetic profile for syk could contribute to the clones’ functional phenotype in response to *Candida albicans*, we challenged each clone with heat-killed yeast and examined the phosphorylation of syk (**Figure 5d-e**). Clones demonstrate greater phosphorylated syk in response to *C. albicans* challenge *in vitro* and clones cluster in their transcriptional pattern closely (**Figure 5e**), suggesting baseline gene expression and open chromatin inform later matured neutrophil behavior to specific stimuli.

**Figure 5.**
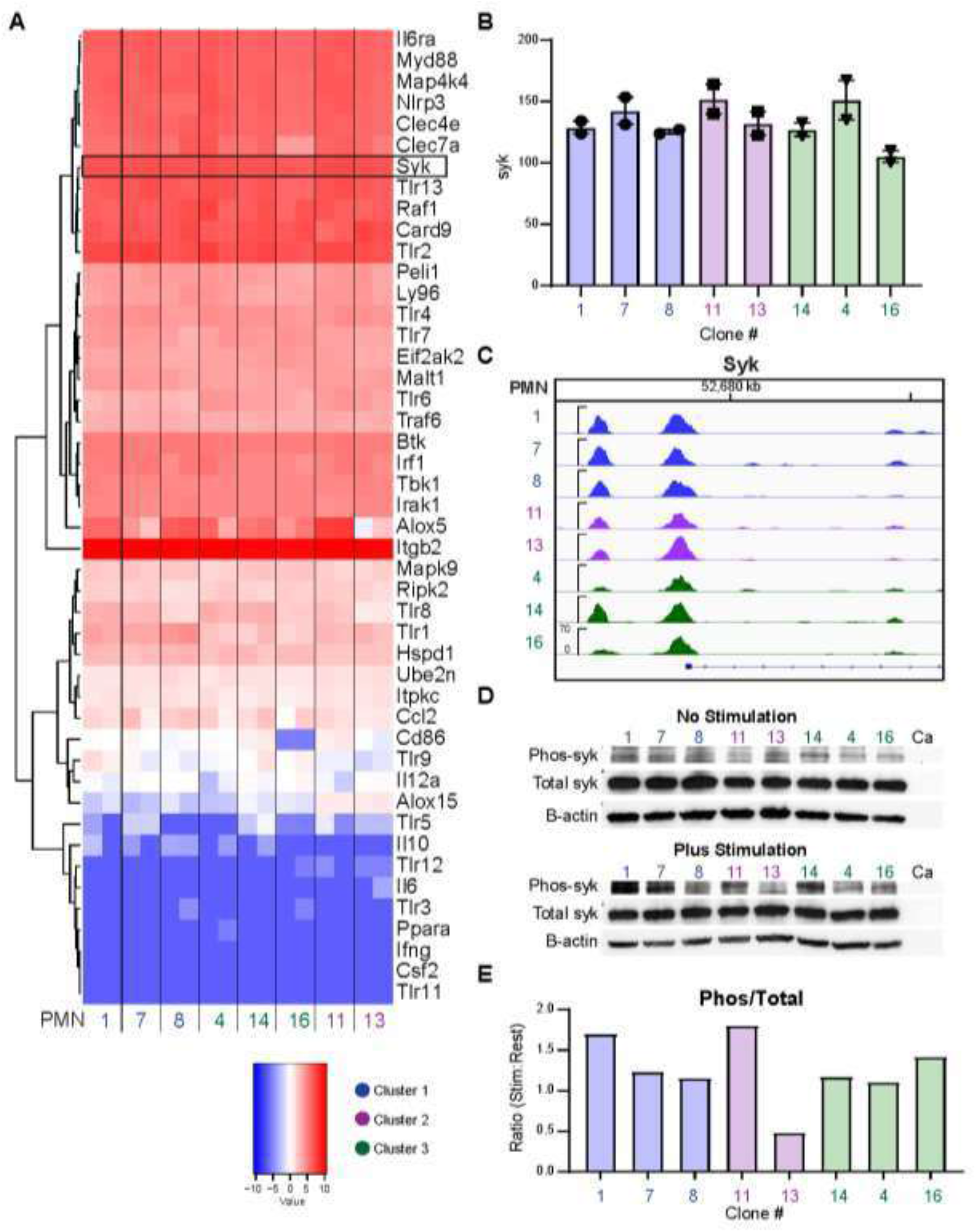
Clone transcriptional profiles indicate areas for functional differences. **(a)** Genes related to neutrophil recognition and response to pathogens were highlighted from the clones’ transcriptional profiling. The heatmap shows the selected genes of interest and clones are shown at replicates. Syk was identified as a pathway of interest and **(b)** the RPKM values for each RNA-seq replicate is shown for each clone. **(c)** The variation in the clones’ open chromatin show relatedness between transcriptional clusters. **(d)** Clones were then challenged with heat-killed *Candida albicans* yeast at MOI 10 and examined for total syk and phosphorylated syk via western blot. **(e)** Quantification of the blots indicates differences between the amounts of phosphorylated syk between clones challenged with *C. albicans*.

### Clonally derived neutrophils demonstrate varied functional responses to *Candida*

Given the distinct clustering of neutrophils by transcription and epigenetic analysis, we sought to determine if clonally derived neutrophils could be functionally distinguished. We challenged sixteen clones with three *Candida spp*. and quantified their effector functions. Neutrophils deploy an arsenal of antimicrobial effector mechanisms to engulf, restrict, and kill pathogens. Here, we tested neutrophil clones for phagocytosis of fungi, reactive oxygen species (ROS) production, fungicidal activity, and neutrophil extracellular trap (NET) release. The prototype *C. albicans* laboratory strain SC5314 induces a potent neutrophil response through ROS and NET production in the hyphal form (Shankar et al., 2020; Hopke et al., 2016). Whereas other *Candida spp*., including *Candida glabrata* and *Candida auris*, produce less-robust neutrophil responses *in vitro* (Negoro et al., 2020). We found that our neutrophils responded uniquely and reproducibly to each of the challenges, and that a vigorous response to one *Candida spp*. did not predict the response to another *Candida spp*. The overall ability of the clones to respond in each assay is illustrated with a spider plot with each spoke corresponding to an individual assay (**Figure 6a**). These plots illustrate overlapping and distinct clonal functions in response to *Candida spp*. and can be utilized as a “fingerprint” to functionally identify each clone.

**Figure 6.**
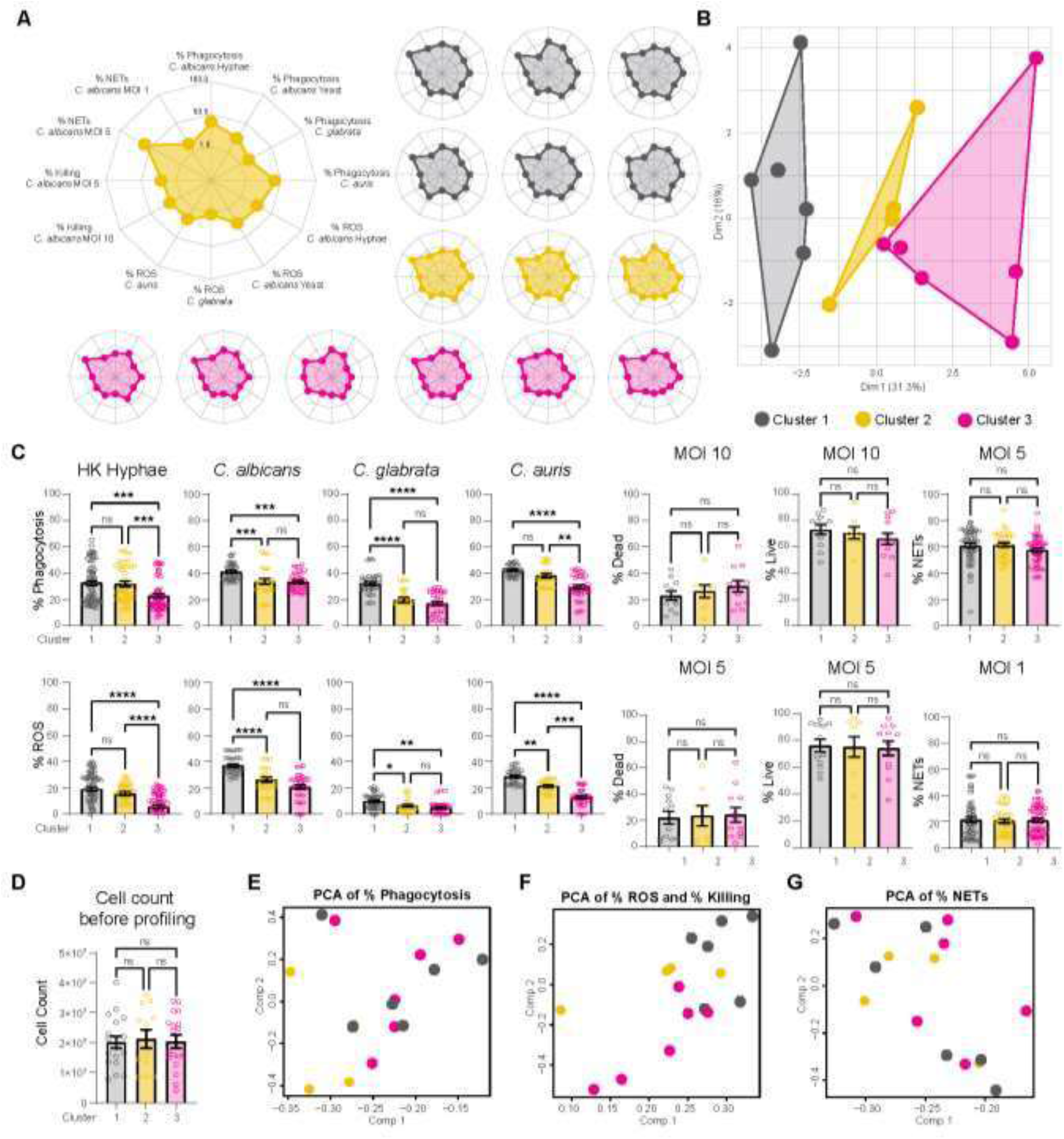
Clonally derived neutrophils demonstrate functional differences in response to fungal challenge. **(a)** Spider plots indicate each clone’s ability to perform known neutrophil effector functions including phagocytosis, reactive oxygen species (ROS) production, killing, and neutrophil extracellular trap (NETs) formation. Each spoke represents a unique response of the clones to three *Candida spp*. **(b)** Clones categorized into three clusters based on functional assays. **(c)** Individual functional assays as shown within the new cluster groups. Individual points represent each clone measured for that assay and error bars indicate SEM. Stats: Kruskall-Wallis with Dunn’s post-test, * p ≤ 0.05, ** p ≤ 0.01, *** p ≤ 0.001, and **** p ≤ 0.001. **(d)** Cell counts are shown from the clones used in the NETs assays as an example of cell division. GMPs were plated as 2 million cells/clones in 10 cm dishes after removal of β-estradiol from the media. Cell counts of the mature neutrophils were taken just before running the NETs assay. Individual points indicate the cell count for a clone for each experiment. Pooled from three NETs experiments, one-way ANOVA with Tukey’s post-test, n.s. not significant. (**e-g**) PCA plots illustrating relatedness between clonal clusters.

Neutrophils control pathogens through phagocytosis and the production of ROS within the phagosome to achieve microbial elimination. We, therefore, tested the ability of clonal neutrophils to phagocytose and produce ROS against three *Candida spp*. Using a flow cytometry-based assay with neutrophils marked by CD11b expression (based on successful neutrophil labeling shown in **Figure 1d)** and heat-killed fungi labeled with AlexaFluor647, we found that clones 1, 2, and 5 were best able to phagocytose and produce ROS consistently between all species of *Candida* while clones 12, 13, 15, and 16 were poorly suited to these tasks (**Suppl. Fig. 6a-d**). However, the clone’s ability to phagocytose *Candida spp*. was not necessarily correlated with the ability to produce ROS when examined by cluster groups (**Suppl. Fig. 6e**). Additionally, clones overall only minimally phagocytosed or produced ROS in response to heat-killed *C. glabrata* which is consistent with previous work using a single wild type clonal GMP cell line(Negoro et al., 2020).

Clones were also tested for their ability to kill a wild-type *C. albicans* strain (SC5314-iRFP) (Hopke et al., 2016) labeled with calcofluor white (CFW). CD11b expressing neutrophils were gated by flow cytometry and examined for the phagocytosis of CFW^+^ fungi. iRFP expression was used to determine the viability of *C. albicans* within the neutrophils. Here, clones 9, 13, and 16 were better able to phagocytose and kill SC5314-iRFP, whereas clones 2, 12, and 14 were able to phagocytose but not kill the pathogen (**Suppl. Fig. 7**). Consistent with the percent phagocytosis of heat-killed *Candida spp*., clones 1, 2, and 5 all showed higher percent phagocytosis scores of live *C. albicans*. Of note, clones 13 and 16 showed poor levels of phagocytosis with heat-killed and live *Candida spp*., but when they did phagocytose live *C. albicans* they were better able to deliver fungicidal activity.

To determine if clonal neutrophils differed in their ability to release NETs, we co-incubated neutrophils with *C. albicans* yeast or hyphae and measured extracellular DNA using Sytox (**Suppl. Fig. 8a-b**). Here, results were normalized to a calcium ionophore positive control (A23187) (Kenny et al., 2017; Sollberger et al., 2018). Specific clonal neutrophils exhibited a robust response to *C. albicans* yeast including clones 1, 10, and 12, which had higher fluorescence marking robust NET release compared to other clones. To ensure the clones were generating true NETs, neutrophils were co-incubated with *C. albicans* hyphae and Sytox green, then counterstained with anti-citrullinated-histone-specific antibody (Ab) Cit-H3 (Hopke and Wheeler, 2017; Negoro et al., 2020). Images of these neutrophils reveal the production of Cit-H3-positive NETs in response to *C. albicans* hyphae indicating true NET formation (**Suppl. Fig. 8c**).

To understand how these effector functions reveal neutrophil heterogeneity, we performed an unbiased clustering approach using the percent scores of all the combined functional assays and found that the clones grouped into three distinct functional clusters (**Figure 6b**). Using these clusters, we pooled the clones for each effector function to examine if parallels existed between the cluster groups and function (**Figure 6c**). The clusters show trends in phagocytosis and ROS production between *Candida spp*. where cluster 1 has a consistently more robust response to fungi than cluster 3. These differences do not appear to be influenced by the rate of cell differentiation *in vitro* as there is no significant difference in cell growth between cluster groups as illustrated in **Figure 6d**. Finally, PCA plots from select effector mechanisms also demonstrate some similar clustering of clones as illustrated by cluster color (**Figure 6e-g**). Together, this suggests that clonal neutrophils can be at least partially defined by their functional responses to *Candida spp*.

### Neutrophil clones demonstrate distinct swarming response to *C. albicans*

To measure more complex, multicellular neutrophil responses that integrate phagocytosis, killing, and other essential functions in response to *C. albicans*, we utilized a recently developed slide-based swarming assay (Hopke et al., 2020). Swarming is an organized behavior of neutrophils in which they initiate and mount effective responses against large pathogen targets, in this case clusters of *C. albicans*, through coordination of neutrophil activities. Here, we simultaneously quantified neutrophil recruitment to the swarm as well as the neutrophil’s function as reflected in the swarm’s ability to restrict *C. albicans* growth. Consistent with our previous *in vitro* findings showing differences between clones, we found that each neutrophil clone swarmed differently in response to *C. albicans*. Following the initiation of a swarming reaction, fungal growth at 16 hrs and the size of the neutrophil swarm at 1 hour displayed differences between clones, with one clone from each functional cluster shown as examples (**Figure 7a**). The total area of fungal growth at the end of the time course was normalized to *C. albicans* growth alone for each clone (**Figure 7b**). Clones demonstrated striking differences in their swarming-specific ability to control the growth of *C. albicans* with clones 1, 7, 14, and 15 showing the highest levels of fungal growth, indicating less overall antifungal capacity. Clones were then pooled based on their cluster groups from Figure 2b and cluster 1 demonstrates better fungal control than either clusters 2 or 3 (**Figures 7c**).

**Figure 7.**
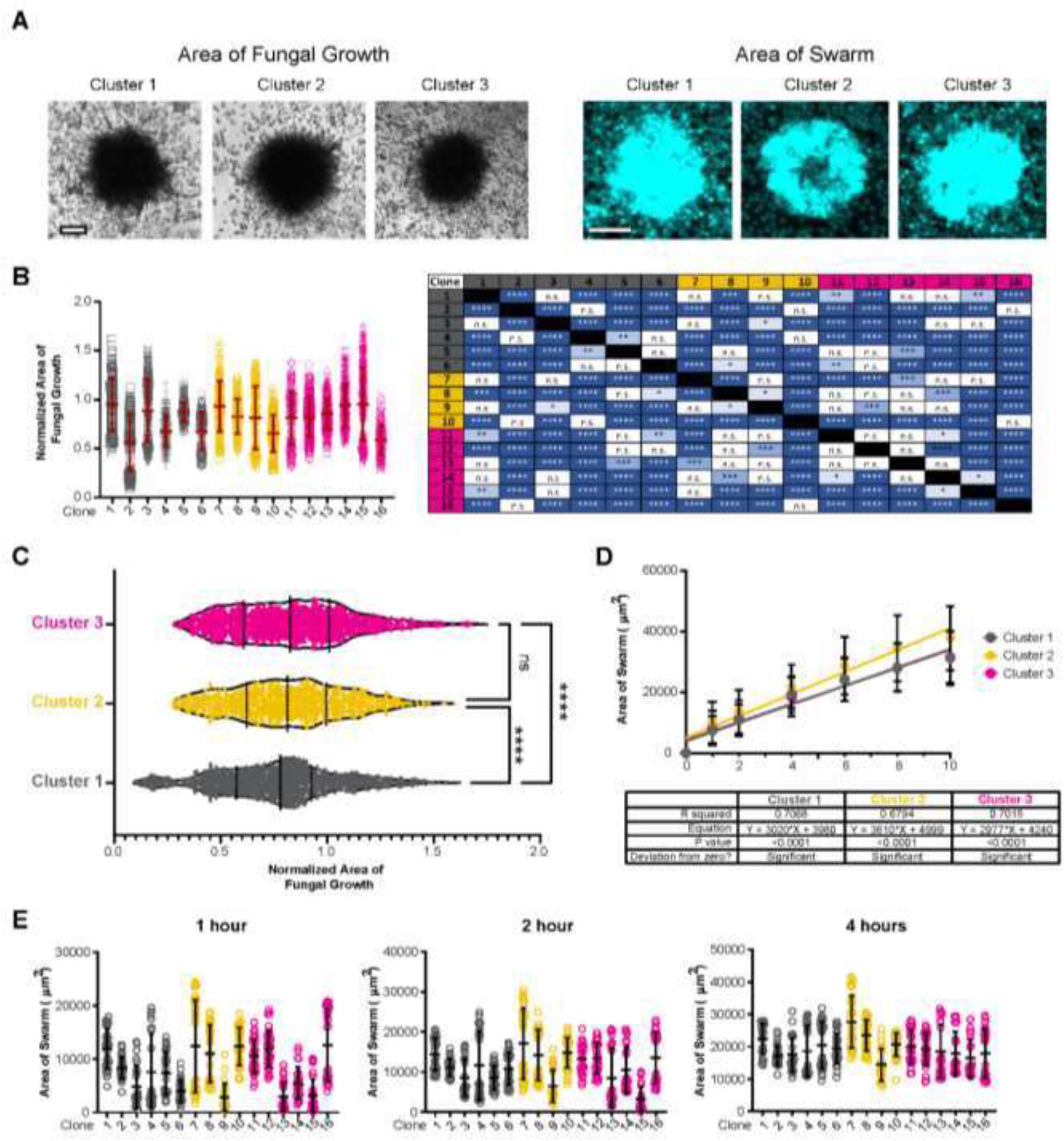
Multi-faceted neutrophil clone swarming identifies unique functional clusters. Clonal neutrophils were added to swarming arrays with clusters of live SC5314-iRFP as the pathogen target. **(a)** Example images of individual targets showing fungal growth (left panels, 16 hours) and Hoechst-stained neutrophils for the area of swarm (right panels, 1 hour) for three different clones, one from each functional cluster. Scale bar, 50 μm. **(b)** The ability of clones to restrict fungal growth was quantified by measuring the total area of fungal growth at 16 hours from the start of the assay. Results were normalized to the area of *C. albicans* that was growing in the absence of neutrophils. Clones were sorted and color-coded by functional cluster. A table showing significant differences in the comparisons between clones is also shown. n = 3 independent experiments with n ≥ 235 swarms quantified per clone. **(c)** Results are also shown as pooled swarms per functional cluster as defined in Figure 6. **(d)** The ability of each clone to form a swarm was also quantified over time (hrs), presented as area of the swarm. The total swarm area was taken for each functional cluster and a line of best fit and R^2^ value calculated for each cluster. **(e)** Area of the swarm is shown for the early timepoints which covers peak neutrophil recruitment. (**d-e**) N = 2 independent experiments with n ≥ 23 swarms quantified per clone. Stats: Kruskall-Wallis with Dunn’s post-test, * p ≤ 0.05, ** p ≤ 0.01, *** p ≤ 0.001, and **** p ≤ 0.001.

Each clone also differed in the robustness of their swarming response, quantified as the area of the swarm over time. Using the same cluster groups, swarming responses for each cluster were quantified (**Figure 7d**). Here, cluster 2 was found to have a more robust swarm than either clusters 1 or 3. In agreement with previous results, swarming initiated quickly and plateaued by 4 hours (**Figure 7e**). The clones did show differences in early dynamics however, with some clones (e.g., clones 1 and 16) showing a more rapid accumulation at early time points (1 and 2 hours) in comparison to others (e.g., clones 6, 9, or 13). Here, clones 7, 8, and 10 demonstrated larger swarm areas by 1 hour (**Figure 7e**) and swarms were fully formed within several hours (**Figure 7d-e** and **Suppl. Fig. 9**). Together, the functional phenotypes of the clonally derived neutrophils support the argument for mature neutrophil heterogeneity.

## DISCUSSION

Here, we describe neutrophil heterogeneity through an examination of neutrophils derived from clonal GMP progenitors. Characterization of clones as immature GMPs and as mature neutrophils demonstrated transcriptional profiling and chromatin accessibility patterns which were maintained through maturation (**Figure 8**). Functional profiling of the mature neutrophils in response to *Candida spp*. illustrated distinct clonal heterogeneity, and our data suggest that neutrophils exist as distinct subpopulations with committed functional phenotypes. Taken together, these data support neutrophil heterogeneity that initiates from the early progenitor stage and that these clonal differences result in phenotypic diversity as illustrated by their clustering profiles (**Figure 8**).

**Figure 8.**
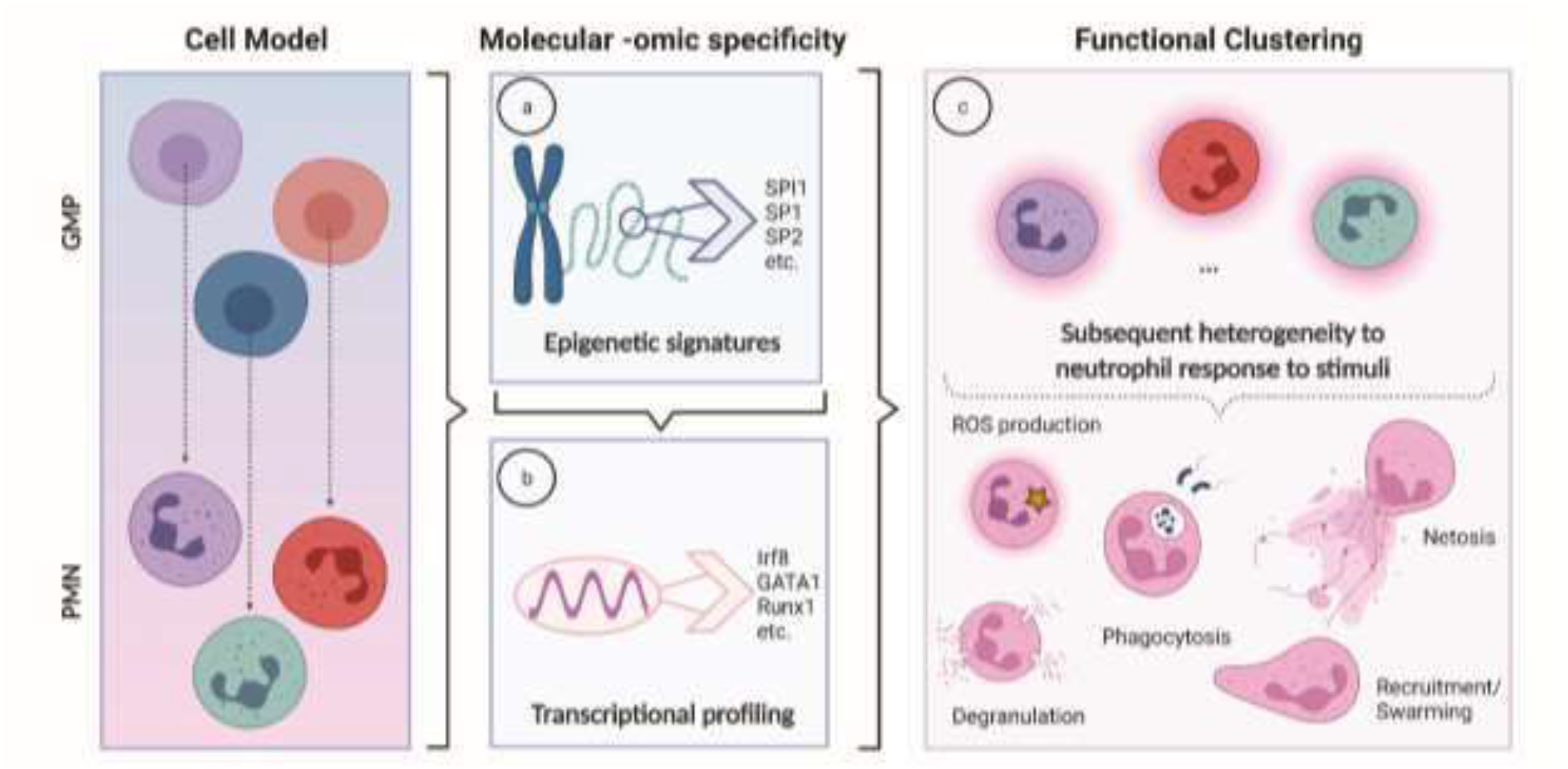
Neutrophil heterogeneity begins at the GMP stage and is preserved into mature neutrophil responses. Illustration of the working model proposed with the clonally derived neutrophil clone system. GMP clones and their corresponding mature neutrophils demonstrate unique epigenetic and transcriptional signatures. Chromatin accessibility at the GMP stage corresponds to later expression of nearby genes at the mature neutrophil stage. The combined epigenetic and transcriptional signatures ultimately result in phenotypically distinct neutrophil clusters in their responses to *Candida*.

Recently, evidence of neutrophil heterogeneity has been demonstrated in homeostasis and disease by transcriptional profiling (Filep and Ariel, 2020; Kwok et al., 2020). While not tied to function, the single-cell transcriptional profiling of murine neutrophils revealed distinct cohorts following a bacterial challenge where unique neutrophil subpopulations were defined in the bone marrow and peripheral blood (Xie et al., 2020). Furthermore, single-cell transcriptional profiling identified two distinct populations of neutrophils that respond in a murine model of pulmonary *Cryptococcus neoformans* infection(Deerhake et al., 2021). These neutrophil populations could be separated by the performance of one or two tasks, namely killing of the fungal pathogen or communicating with other immune cells at the site through cytokine production (Deerhake et al., 2021). Notably, these differences in neutrophils could also represent how tissue specificity impacts neutrophil heterogeneity and future work may address how neutrophil populations change in tissues in response to infection. Finally, functional profiling of human healthy donor neutrophils has also demonstrated potential links to gene expression from bulk RNA-sequencing (Maskarinec et al., 2022).

To explore and define heterogeneity beyond functional profiling, we performed transcriptional and epigenetic analyses. The data link functional clustering of neutrophil clones to their transcriptional and epigenetic grouping. RNA-seq based clusters define basic neutrophil heterogeneity under homeostatic conditions, further investigation will be required to map the evolution of neutrophil phenotypes in response to *Candida spp*. Additionally, more work will be required to define changes in functional clusters following inflammatory stimuli or infection. We identified several gene pathways that exhibit delayed expression, but which also demonstrated the priming of specific genes as evidenced by enhancer chromatin accessibility at the GMP stage. Recently it was demonstrated that trained immunity could be passed down from mother to offspring and that these progeny exhibit altered progenitor cell development at the transcriptional and epigenetic level (Katzmarski et al., 2021).

Our transcriptional data demonstrate that conditionally immortalized GMP clones mature into neutrophils which are committed to a specific functional capacity when challenged with *Candida spp*. To determine if heterogenous clustering existed at a function level, we applied functional clustering encompassing a large array of metrics including fungicidal killing, the generation of reactive oxygen species (ROS), phagocytosis, neutrophil extracellular trap (NET) production, and multi-functional swarming. Unbiased clustering of these functional metrics confirmed several distinct clusters. These clusters were also present in a multi-faceted swarming assay in which neutrophil recruitment and control of *C. albicans* could be examined in a single experiment (Hopke et al., 2020). Interestingly, neutrophil control of fungal growth did not correlate directly with the rate of neutrophil accumulation when examined in swarming assays. Specific neutrophil clones accrued quickly but controlled fungal growth poorly, indicating that effective fungal clearance requires multiple effector mechanisms besides response and recruitment to the site of infection. Neutrophils need to achieve an optimal balance of rapid response and effective antimicrobial function, something which our clonal phenotyping signatures highlight. These results suggest that clone-specific pre-positioning exists and supports the hypothesis that defined neutrophil heterogeneity exists at homeostasis at least from the GMP stage.

Our experimental model system involves the use of a retrovirally-delivered estrogen receptor homeobox B8 (ER-HoxB8) fusion protein (Wang et al., 2006; Sykes and Kamps, 2001) to circumvent the limitations of the short neutrophil life span of <24 hours and the lack of genetic tractability. The ER-Hoxb8 system has now been widely adopted and proven a reliable and advantageous tool for the study of innate immune cells including monocytes (Accarias et al., 2020), dendritic cells (Cabron et al., 2018), eosinophils (Ma et al., 2020), basophils (Gurzeler et al., 2013), and neutrophils (Chu et al., 2018; Saul et al., 2019; Orosz et al., 2020). Successfully transduced GMPs were expanded to sufficient cell numbers for functional phenotyping. The conditional immortalization system relies on viral transduction and random integration of the ER-HoxB8 transgene. The random retroviral insertion into the GMP genome raises the potential for non-specific alternation of up or downstream gene expression which could influence GMP phenotype, although we feel that likelihood of this phenomenon is low given the non-random clustering of clones at a functional, transcriptional, and epigenetic level. In future studies, we will circumvent this limitation through the generation of a transgenic mouse line in which the ER-Hoxb8 transgene is expressed from a single defined locus. Moreover, while this limitation may be at play, this study is a genuine proof-of-concept of a genetically tractable system permitting for the first the examination of heterogeneity in neutrophils and, in the future, other short-lived hematopoietic cell types.

Our data demonstrate the existence of heterogeneous neutrophil precursors at the GMP stage in bone marrow (**Figure 8**). The identification of marked open chromatin resulting in delayed expression of set pathways in mature neutrophils raises the potential of an epigenetic phenomenon as the putative mechanism for the generation of neutrophil heterogeneity. We demonstrate that these GMPs appear to be predestined to a specific functional cluster in response to *Candida spp. in vitro*. Whether an epigenetic predetermined state occurs at the GMP stage or via an earlier stem cell is yet to be defined. In addition, the rules governing these predestined heterogenous functional clusters and if they are fluid or adaptable in the setting of inflammation or infection will require additional investigation. Finally, these results begin to define the molecular mechanisms required to potentially augment anti-fungal responses in the neutrophil compartment, permitting the development of needed immunotherapies for those at the highest risk for invasive fungal infections.

## METHODS

### CD45.1^STEM^ single-cell clone construction and maintenance

Single-cell clones were generated from 6-8-week-old female C57Bl/6 CD45.1^STEM^ mice (Mercier et al., 2016) housed in the specific-pathogen-free animal facility at Massachusetts General Hospital (MGH) in accordance with the MGH Institutional Animal Care and Use Committee (IACUC; #2017N000058). Granulocyte monocyte progenitors (GMPs) were isolated for transduction of the estrogen receptor-homeobox B8 (ER-HoxB8) gene fusion as described previously (Wang et al., 2006). Briefly, bone marrow from the femurs and tibias was harvested and layered over Ficoll-Paque PLUS (GE Healthcare). Cells were harvested from the interface, washed with FACS buffer (phosphate-buffered saline (PBS), 2% fetal bovine serum (FBS), and 1 mM EDTA), and brought to 2×10^6^ cells/ml in cytokine stimulation media containing: RPMI 1640 with 2 mM L-glutamine (Corning, Manassas, VA), 10% heat-inactivated fetal bovine serum (Life Technologies, Grand Island, NY) 1% penicillin-streptomycin (Thermo Fisher Scientific, Waltham, MA), stem cell factor (SCF), interleukin-3 and interleukin-6 (each at 10 ng/ml, Peprotech, Rocky Hill, NJ). SCF was prepared as described previously (Negoro et al., 2020). Briefly, SCF-overexpressing Chinese hamster ovary cells were cultured, the SCF-conditional media was harvested, tested for activity, filtered, and stored at –80°C (Negoro et al., 2020; Sykes et al., 2016; Sykes and Kamps, 2001). The harvested cells expanded for 36-48 hours, brought to 5×10^5^ cells/ml, and transduced with the ER-HoxB8 construct (Wang et al., 2006). Retroviral infection was done in human plasma fibronectin (10 μg/ml, Sigma Aldrich) coated 12-well plates with Polybrene (24 μg/ml; Millipore, Burlington, MA) and spinoculated at 1000xg for 90 min at 22°C. GMPs with the ER-HoxB8 construct successfully integrated were selected for with 2 mg/ml geneticin (G418; Thermo Fisher Scientific, Waltham, MA). Single-cell clones were sorted into a 96-well U-bottom plate via a BD FACSAria flow cytometer (BD Biosciences) at the Center for Regenerative Medicine (CRM) Flow Cytometry Core Facility (MGH, Boston, MA) in β-estradiol (0.5 μM, Sigma Aldrich) containing media. Sixteen single cell sorted clones were selected from the 96 well plate and used in this study.

GMPs were maintained in culture in the presence of SCF and β-estradiol which permits nuclear translocation of the ER-HoxB8 fusion protein, resulting in the conditional immortalization of the GMP (Wang et al., 2006; Negoro et al., 2020). Removal of β-estradiol from the media restores differentiation of the GMP into mature neutrophils (Wang et al., 2006; Negoro et al., 2020). To remove β-estradiol completely, GMPs were washed twice with phosphate-buffered saline (PBS; Corning, Manassas, VA) prior to seeding cells in complete RPMI containing SCF. All cells were grown in a humidified incubator at 37°C in the presence of 5% CO_2_.

### Maturation of single cell clones

To ensure the selected GMP clones developed into mature neutrophils, GMPs were removed from β-estradiol and allowed to mature for four days. Cells were washed with PBS, brought to 5×10^5^ cells/ml, and labelled with FITC-labeled monoclonal antibody anti-Mac-1 (CD11b) from BioLegend (San Diego, CA), CD117 (BioLegend, San Diego, CA), Ly6G (BioLegend, San Diego, CA), and F4/80 (BioLegend, San Diego, CA) for 30 min at 4°C. Cells were washed with FACs buffer and run on a FACsCalibur or BDCelesta flow cytometer.

### Cell surface labelling and flow cytometry

Clones were labelled for cell surface markers as described previously (Xie et al., 2020) with the following changes. Briefly, clones were removed from β-estradiol containing media four days prior to use, brought to a concentration of 5×10^6^ cell/ml, and fixed in 10% formalin for 30 min. at room temperature. Cells were washed twice in FACs buffer containing 2% FBS and incubated in the dark with panel G1-4 antibodies: APC-conjugated CD117, FITC-conjugated CD34, BV421-conjugated Ly6G, and PE-conjugated CXCR2 for 30 min. at 4°C or, cells were labelled with panel G5a-c antibodies as described. Cells were first incubated for 30 min. at 4°C in the dark with anti-IFIT1/p65 antibody (Sigma-Aldrich), washed with FACs buffer, and then incubated with a master mix solution containing APC-conjugated CD45.1, APC/Cy7-conjugated CD11b, PE-conjugated Ly6G, BV421-conjugated CXCR4, and with the secondary goat anti-rabbit-Alexa Fluor 488 antibody (Invitrogen) for 30 min. at 4°C in the dark (see also Table 1). Cells fixed for supplementary figure flow panels. Otherwise, cells were labelled for 30 min. at 4°C, washed with FACs buffer, and analyzed on a BDCelesta (BD Biosciences) flow cytometer. GMPs were also labelled for progenitor markers using the panel in Table 1. (Challen et al., 2009) All flow cytometry analysis was completed with FlowJo software (v10.7.1 for Windows, Becton, Dickinson and Company, 2019).

### Bulk RNA isolation, sequencing, and analysis

Clonal naïve GMP and neutrophil total RNA was extracted from cells at 5×10^6^ cells/ml using the Qiagen RNeasy Plus Mini Kit (Qiagen) with DNaseI treatment (RNase free DNase Set, Qiagen). RNA quality was confirmed by RNA Tapestration and quantified with Qbit. All samples showed RNA quality of >7.5 and RNA-seq libraries were prepared. Sequencing was performed on Illumina HiSeq 2500 instrument, resulting in approximately 30 million of 50 bp reads per sample. Sequencing reads were mapped in a splice-aware fashion to the mouse reference transcriptome (mm9 assembly) using STAR (Dobin et al., 2013). Read counts over transcripts were calculated using HTSeq (Anders et al., 2015) based on the Ensembl annotation for NCBI37/mm9 assembly. For differential expression analysis, we used the EdgeR method (Robinson et al., 2010) and classified genes as differentially expressed based on the cutoffs of 2-fold change in expression value and false discovery rates (FDR) below 0.05. For clone-to-clone comparisons, the cutoff of 2-fold change in expression value was used. A minor batch effect observed among RNA-seq samples of neutrophil clones was corrected using the Combat method (Johnson et al., 2007) implemented in the SVM R package (Leek et al., 2012).

### ATAC-Seq preparation and analysis

Naïve GMP and neutrophil paired clones were prepared for ATAC-seq as previously described (Buenrostro et al., 2015). Briefly, cells were brought to 5×10^5^ cells/ml and incubated with transposition reaction mix (NEBNext High-Fidelity 2X PCR master mix, NEB) for 30 min. followed by PCR for 18 cycles. ATAC-seq samples were not size selected here. ATAC-seq sequencing was performed on Illumina HiSeq 2500 instrument, resulting in approximately 15-20 million of paired-end 50 bp reads per sample. Reads were mapped to mm9 mouse reference genome using BWA (Li and Durbin, 2009). Fragments with both ends unambiguously mapped to the genome that were longer than 100 bp were used in further analysis. Hotspot2 (John et al., 2011) was used to detect significant peaks with FDR cutoff of 0.05. The resulting peak regions were analyzed for changes in read density between conditions. For the analysis of overlap between peak regions, we used the cutoff of 50% reciprocal overlap between the two compared regions. For the analysis of differential chromatin accessibility between groups of replicate samples, DiffBind R package was used (Ross-Innes et al., 2012; Stark and Brown, 8AD).

### *Candida* strains and growth conditions

Wild type *C. albicans* was purchased from the American Type Culture Collection (ATCC MYA-2876; ATCC, Manassas, VA) while the far-red fluorescent (SC5314-iRFP) *C. albicans* strain was a kind gift from Robert Wheeler (University of Maine, Orono, ME) (Hopke et al., 2016). Strains were streaked from frozen stock onto yeast extract peptone dextrose (YPD; 1% yeast extract, 2% peptone, 2% dextrose, Sigma-Aldrich) agar and left overnight at 37°C. Single colonies were inoculated to YPD broth and grown overnight at 30°C with shaking. *C. glabrata* (ATCC 2001; ATCC, Manassas, VA) and *C. auris* (sequence confirmed clinical isolate, South American clade; MGH microbiology lab, Boston, MA) cultures were inoculated from frozen culture directly to YPD broth and incubated overnight at 30°C with shaking. After incubation, cultures were washed twice with PBS, counted with a Luna automated cell counter (Logos, Biosystems, Annandale, VA), and resuspended in PBS at the desired inoculum. Where noted, *Candida spp*. were heat-killed on a 95°C heating block for 20 min., washed once with PBS, then labelled with AlexaFluor647 (Life Technologies, Eugene, OR) for 30 min with shaking in the dark, washed twice with PBS, and brought to the desired inoculum.

### Western blot

Clones were collected at day 4 out of β-estradiol and brought to 2.5×10^6^ cells/ml in cRPMI. Neutrophils were co-incubated with heat-killed *C. albicans* at MOI 10 for 40 minutes at 37°C, 5% CO2. Samples were lysed in Laemmli sample buffer (Bio-Rad, Hercules, CA) containing protease inhibitors (cOmplete mini; Roche Diagnostics, Indianapolis, IN), reducing agent (NuPAGE sample reducing agent; Thermo Fisher Scientific), and phosphatase inhibitors (sodium orthovanadate; New England Biolabs, Ipswich, MA). Proteins were resolved by SDS-PAGE, transferred onto a polyvinylidene difluoride membrane, and blocked with Tris-buffered saline–1% Tween (TBS-T)–5% nonfat milk. Total Syk protein was detected with rabbit MAb D3Z1E anti-Syk (Cell Signaling, Danvers, MA) (1:1,000) and phosphorylated Syk was detected with rabbit Mab 2710 anti-Phospho-Syk (Tyr525/526) (Cell Signaling, Danvers, MA) (1:1000). The blots were subsequently reacted with mouse MAb AC-15 anti-β-actin (Sigma, St, Louis, MO) (1:200,000) to verify equivalent loading levels.

### Phagocytosis and ROS assessment

Neutrophil clones were collected at day 4 out of β-estradiol and brought to 5×10^5^ cells per condition. Neutrophils were co-incubated with heat-killed *Candida spp*. at MOI 10 for *C. albicans* hyphae for 2 hour and MOI 3 for *C. albicans* yeast, *C. glabrata*, and *C. auris* for 5 hours. Dihydrorhodamine-123 (DHR-123, Life Technologies, Eugene, OR) was added to each condition at a final concentration of 1 μM. Cells were co-incubated at 37°C with CO_2_ before fixation in 1% formalin for 30 min, then washed and resuspended in FACs buffer prior to labelling and flow. Cells were labelled with CD11b-PE (clone M1/70, Biolegend, San Diego, CA) on ice for 30 min prior to flow. Samples were run in triplicate on a FACsCalibur using the CellQuest software in a 96 well round bottom plate. Neutrophils were gated using CD11b surface marker expression and the percent phagocytosis of *Candida spp*. or percent fluorescence of the DHR-123 was analyzed with FlowJo 10 software. The average background of DHR-123 fluorescence was subtracted from each *Candida* condition.

### Fungal killing assay

Clones were brought to 5×10^5^ cells per condition and co-incubated with live SC5314-iRFP at a MOI 10 or MOI 5 for 2 hrs at 37°C. Samples were fixed in 1% formalin for 30 min., washed with FACs buffer, and stored at 4°C in the dark. Neutrophils were labelled with CD11b-FITC (clone M1/70, Biolegend, San Diego, CA) and *C. albicans* was labelled with calcofluor white (CFW) on ice for 30 min. prior to being run on the LSRII flow cytometer at the CRM Flow Cytometry Core Facility (MGH, Boston, MA). Neutrophils containing *Candida* were selected by CD11b-FITC and then examined for CFW positive and iRFP expression from the fungi with FlowJo 10 software. Percent live and percent dead fungus was calculated from the total number of CFW labelled cells.

### Neutrophil extracellular trap (NET) formation

Neutrophils were plated at 1×10^5^ cells per well in complete RPMI in a 96 well plate after 4 days out of β-estradiol. *C. albicans* yeast at MOI 1 or *C. albicans* hyphae at roughly MOI 5 were co-incubated with Sytox green (final concentration 1 μM) (Life Technologies, Eugene, OR) for 5 hours at 37°C with 5% CO_2_. After incubation, the plate was read on a SpectraMax i3x for Sytox fluorescence (excitation 500 nm/emission 528 nm) to detect extracellular DNA. The average background fluorescence measured from resting neutrophils with Sytox was subtracted from each condition and results were expressed as arbitrary fluorescence units. The calcium ionophore A23187 (Sigma-Aldrich) was used as a positive control at a final concentration of 5 μM(Sollberger et al., 2018; Kenny et al., 2017) and fluorescent reads were used to normalize the data. To determine if extracellular DNA was associated with NETs, clones were co-incubated with *C. albicans* hyphae for 3 hours and then labelled with anti-histone H3 citrulline R2+R8+R17 (Abcam, Cambridge, MA; 0.014 mg/ml) followed by donkey anti-rabbit IgG Cy3 (Jackson Immunoresearch, West Grove, PA; 0.0075 mg/ml) secondary antibody, as previously described (Hopke and Wheeler, 2017). Samples were imaged at x40 on a Nikon Ti-E microscope to demonstrate NET release for each clone.

### Array printing and image analysis for neutrophil swarming

Swarming assays were conducted as previously described(Hopke et al., 2020). Briefly, a solution of 0.1% poly-l-lysine (Sigma-Aldrich) was printed via a microarray printing platform (Picospotter, PolyPico, Galway, Ireland) as 100 μm diameter spots for *C. albicans* adherence. For experiments, 8 × 8 arrays in a 16-well format on ultra-clean glass slides (Fisher Scientific) were prepared and screened for accuracy. Slides were left at 40°C for 2 hours on a heated block to dry then left at room temperature. All imaging was conducted using a fully automated Nikon TiE microscope. Time-lapse imaging was conducted using a ×10 Plan Fluor Ph1 DLL (NA= 0.3) lens and endpoint images were taken with a ×2 Plan Apo (NA= 0.10) lens. Confocal imaging was conducted using a ×40 Plan Fluor (NA= 0.75) lens. Swarming targets were selected and saved using the multipoint function in NIS elements prior to loading of neutrophils. Five hundred thousand clonal neutrophils were added to each well. All selected points were optimized using the Nikon Perfect Focus system. Area analysis was performed manually by outlining the swarms or areas of fungal growth in the NIS-elements (v4.00.12; Nikon Inc.) or FIJI (FIJI is just ImageJ v2.0.0-rc-59/1.52p, NIH) software. For area of swarm, only the neutrophil swarm identified by the Hoechst-stained cells was measured. Areas of fungal growth were measured through brightfield and fluorescent channels.

### Software and statistical reproducibility

All flow cytometry analysis was completed with FlowJo software (v10.7.1 for Windows, Becton, Dickinson and Company, 2019). Data were tested for normality using a D’Agostino–Pearson omnibus normality test. Normally distributed data were analyzed with Student’s T test or one-way analysis of variance with Tukey’s post-test. Non-normally distributed data were analyzed with Kruskal–Wallis with Dunn’s post-test where appropriate. Statistical significance was considered for p < 0.05; exact p values are provided in the relevant figure legends. All statistics were conducted using the GraphPad Prism (v8.2.1 for Windows, La Jolla, CA).

## Acknowledgements

The authors would like to thank the Center for Regenerative Medicine Flow Cytometry Core Facility at MGH, with special gratitude to M. Handley, for technical assistance and insight. Additional experimental assistance was provided by S. Mittiga, A. Crossen, D. Vargas, J. Reedy, and H. Brown. Helpful discussions and comments were provided by B. Ward and past and present Vyas, Mansour, and Sykes lab members.

## Author Contributions

Project initiation, A.K.S., S.X., D.B.S., M.K.M.; Experimental design and operation, A.K.S., A.H., S.X., M.K.M.; Experimental workflow assistance, A.H., A.V., N.J.A., K.T., D.E.A., C.R.; Functional data analysis, A.K.S., A.H.; Sequencing data analysis, M.C., R.S.; Provided advice and discussion, A.H., N.A., D.S., D.B.S., D.I., R.S., M.K.M.; Writing – original draft, A.K.S., A.H., M.C., R.S., M.K.M.; Supervision and funding acquisition, M.K.M. All authors participated in manuscript editing and approved the final manuscript.

## Data and Materials Availability

Supporting data for this study is available from Dryad (https://doi.org/10.5061/dryad.q83bk3jm) and from the corresponding author upon request. All sequencing data generated in this study have been deposited at NCBI’s Gene Expression Omnibus (GEO) repository and are accessible through GEO Series accession number GSE188683 and GSE188682.

---

**Supplementary Figure 1.**
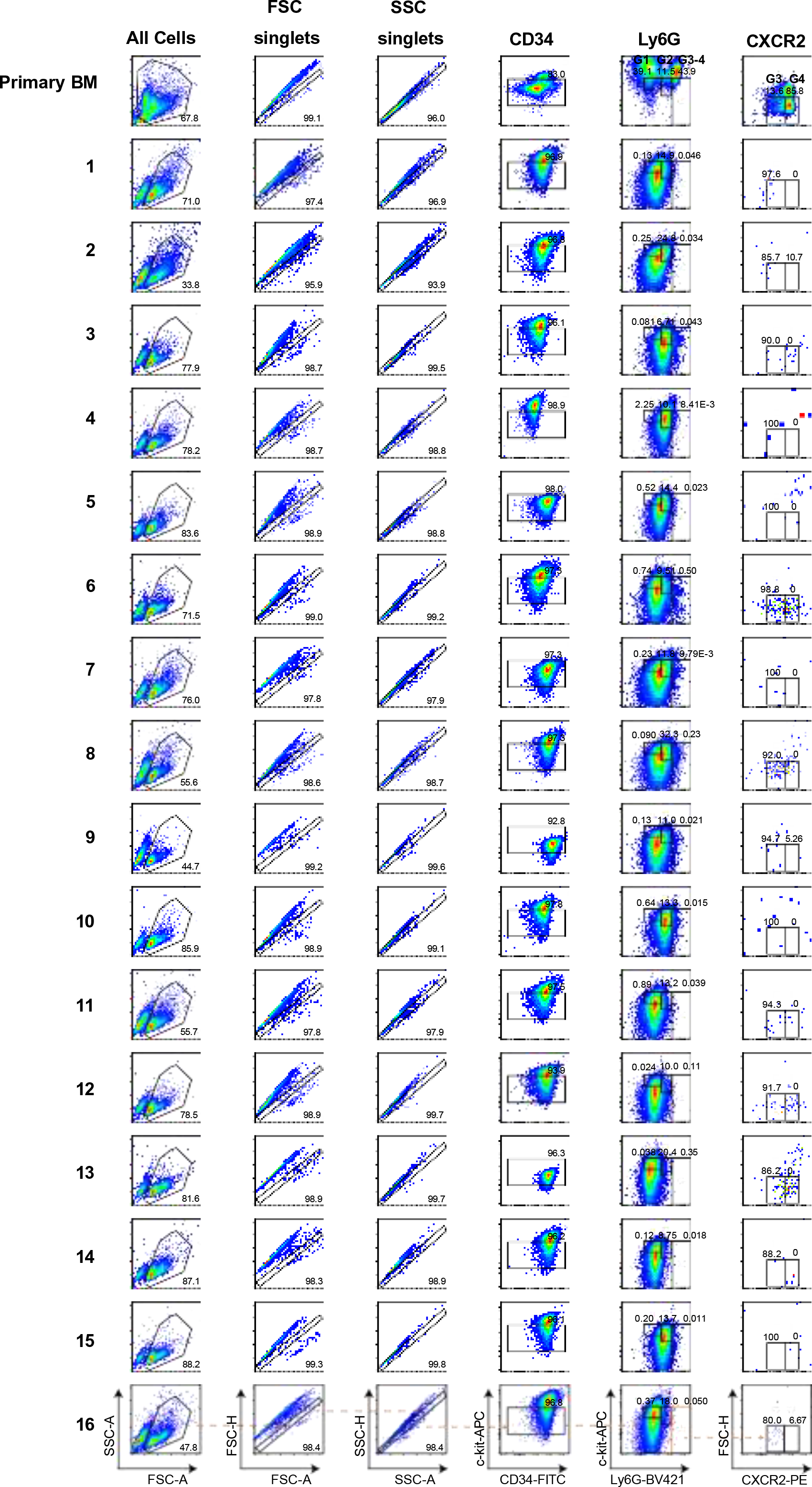
Gating scheme for GMP markers. GMP clones were counted and labelled for cell surface markers to indicate their maturation status. Clones were compared with bone marrow harvested from a wildtype mouse which was used to make the gating pattern for the progenitor pattern.

**Supplementary Figure 2.**
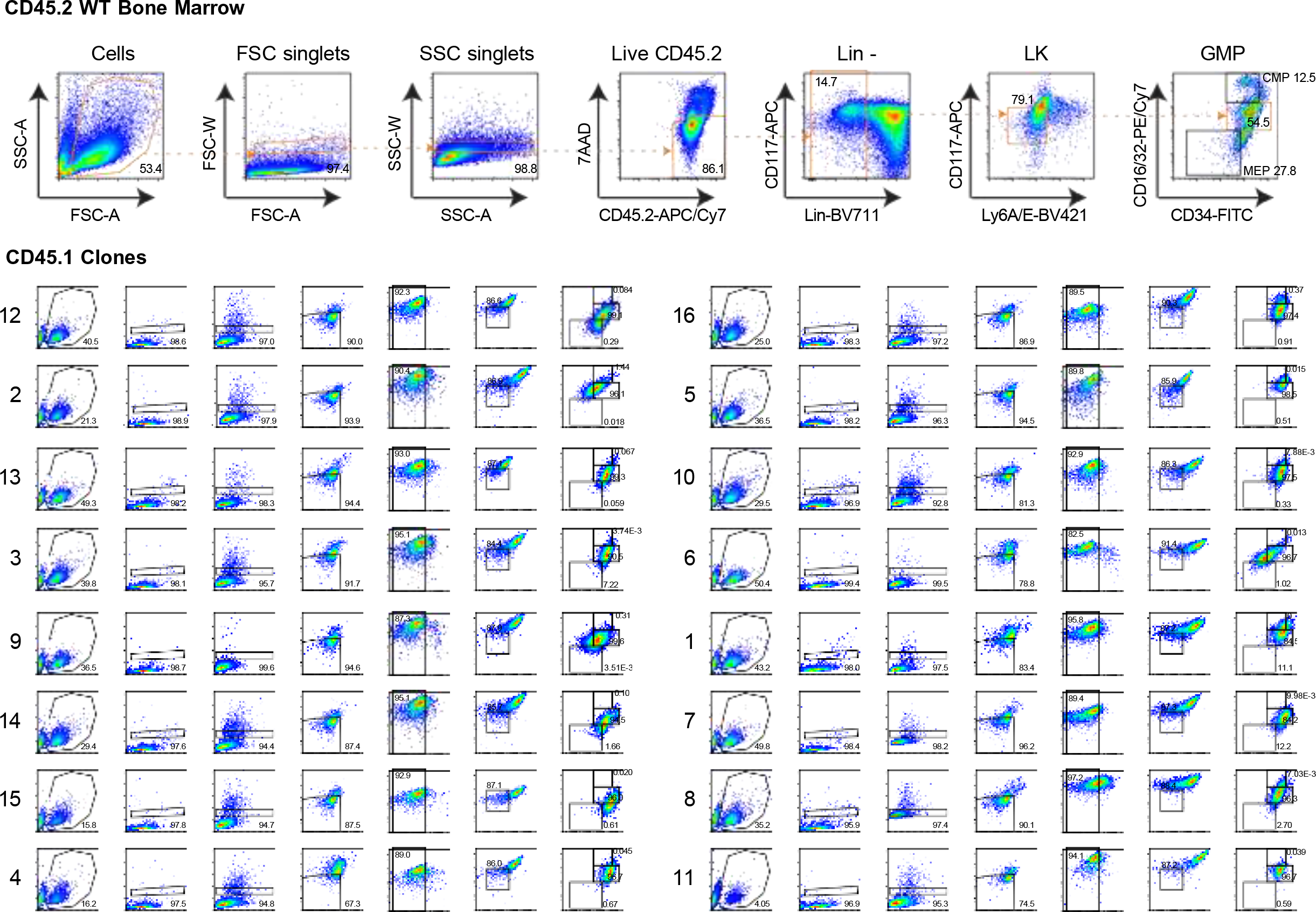
Gating scheme for immature neutrophil markers. GMP clones were counted, fixed, and labelled for cell surface markers to indicate their maturation status. Clones were compared with bone marrow harvested from a wildtype mouse which was used to make the gating pattern for G1-G4 immature to mature neutrophils.

**Supplementary Figure 3.**
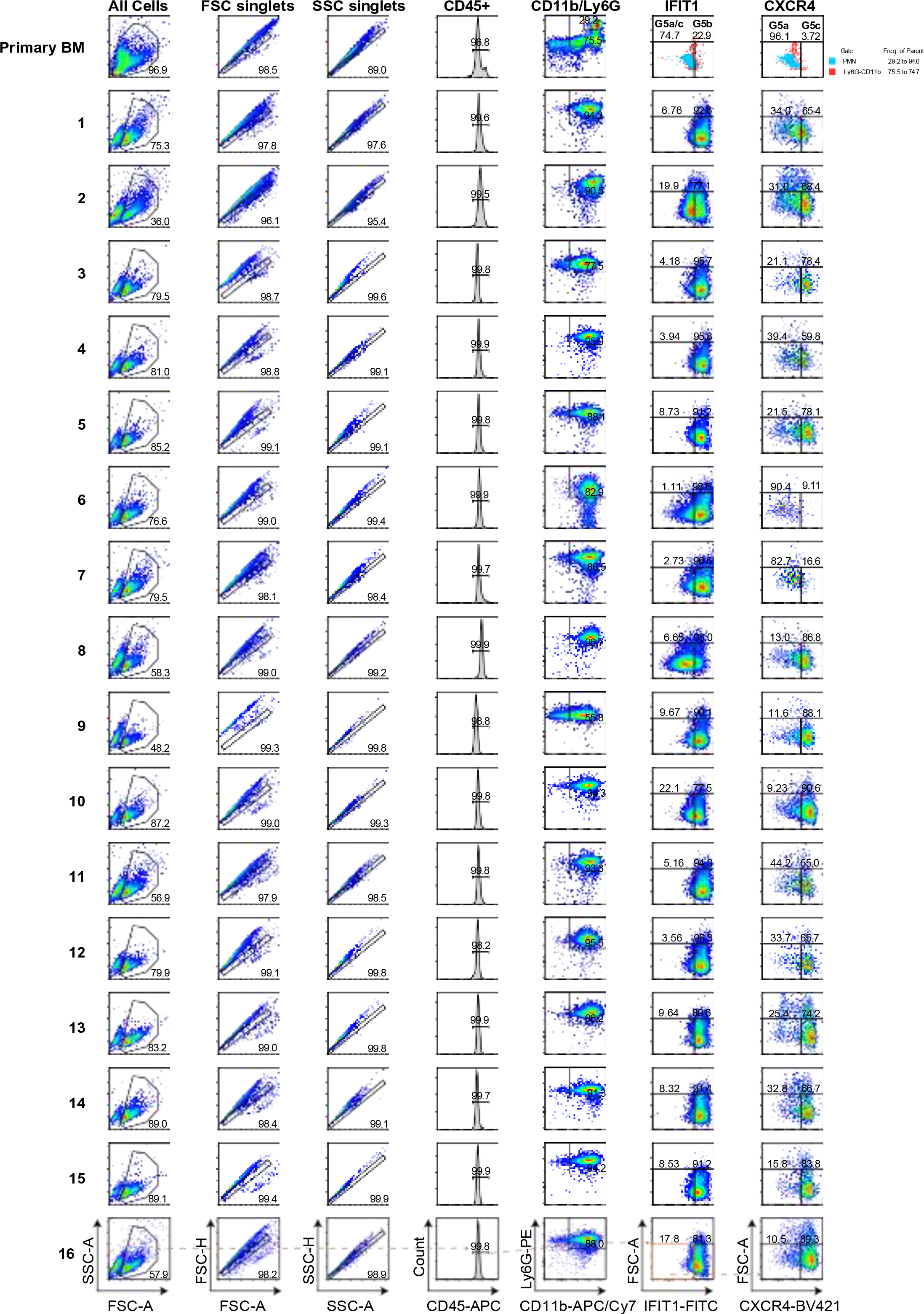
Gating scheme for mature neutrophil markers. GMP clones were removed from estradiol and allowed to mature for 4 days. Then neutrophils were counted, fixed, and labelled for cell surface markers to indicate their maturation status. Clones were compared with bone marrow harvested from a wildtype mouse which was used to make the gating pattern for G5a-c mature neutrophils.

**Supplementary Figure 4.**
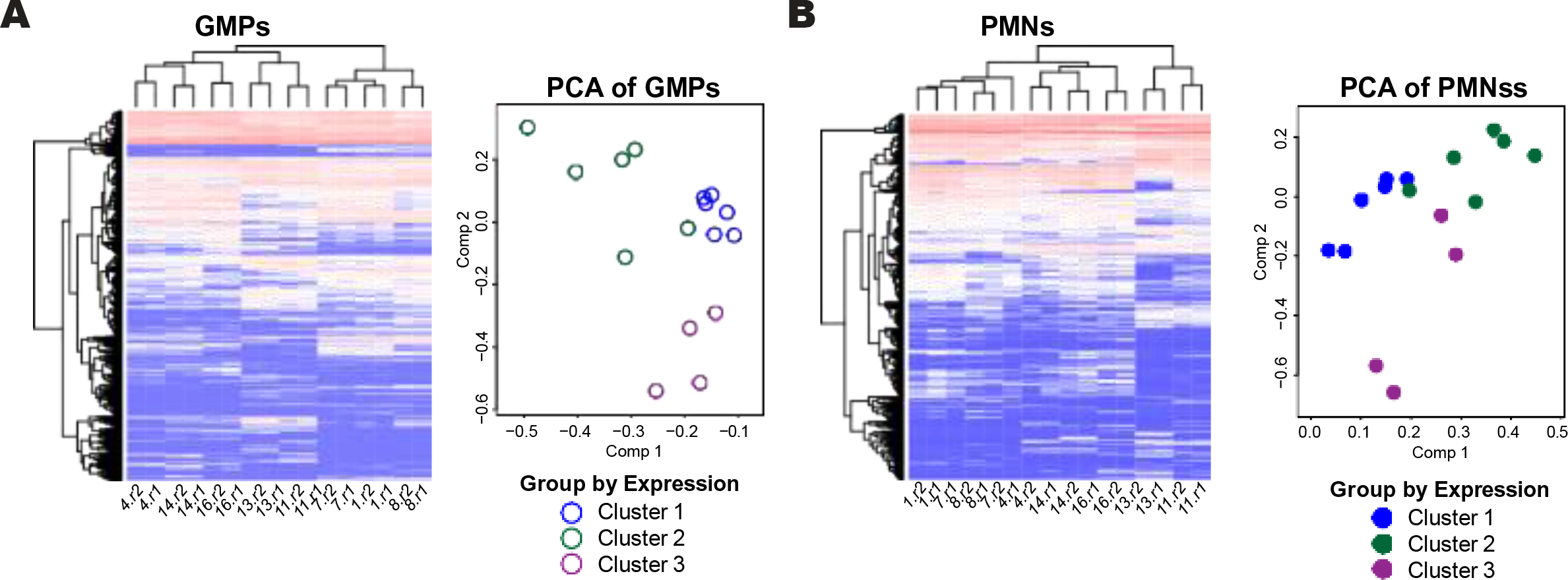
RNA-sequencing results in secondary cluster pattern. **(a)** Heatmap (log2(RPKM)) and PCA plots with 860 DEGs in the union GMPs and **(b)** 523 DEGs in the union for neutrophils. Colors indicate clustering method based on gene expression.

**Supplementary Figure 5.**
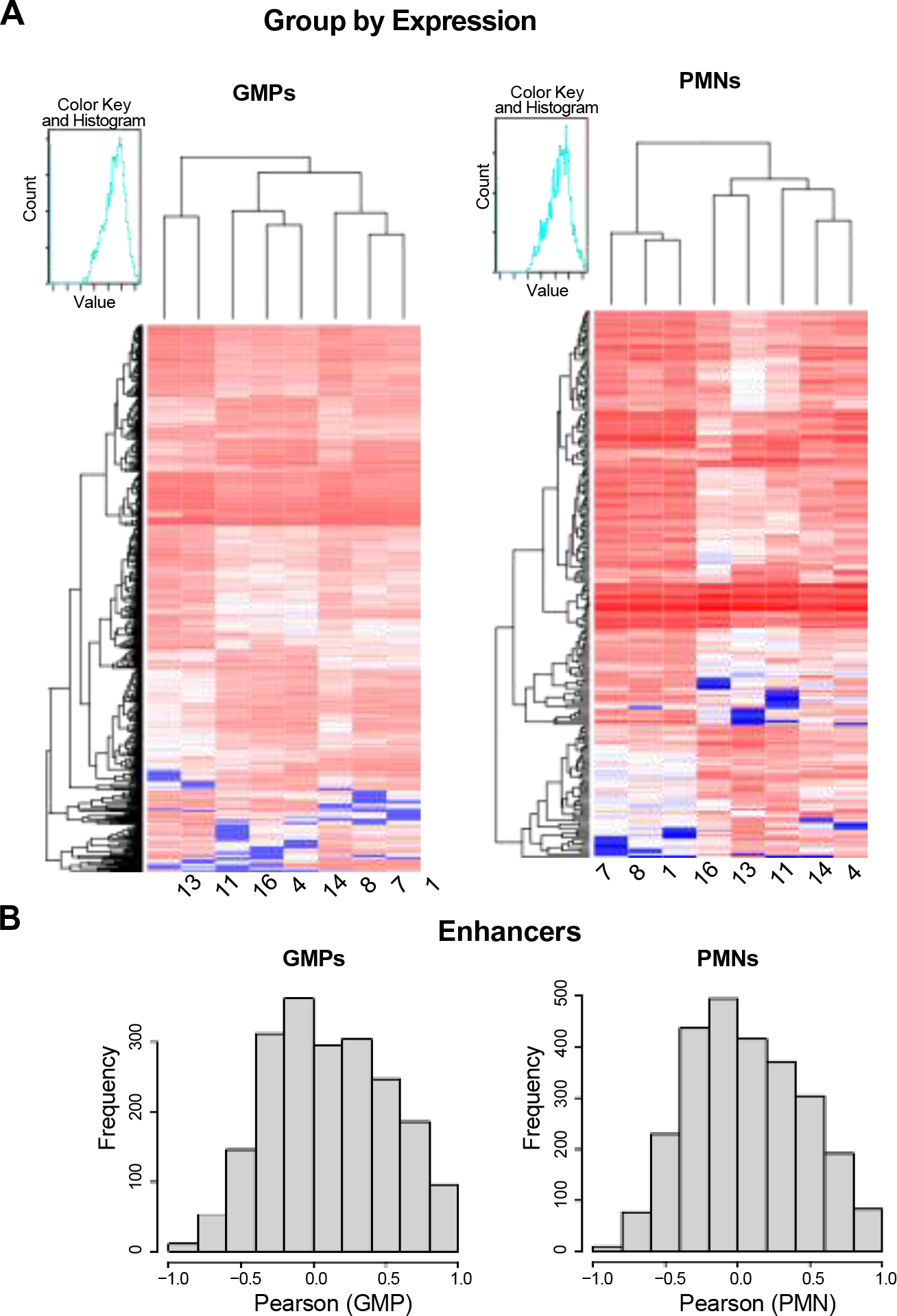
Chromatin accessibility demonstrates modest correlation with clone gene expression. **(a)** Heatmaps corresponding to the PCA plots in **Figure 5e-f** based on expression grouping. **(b)** Clones were compared one-by-one within GMPs and neutrophils to obtain the union of DEGs from RNA-sequencing and the union of DARs from ATAC-Seq. For the union of DEGs, we found corresponding promotors and enhancers DARs. The Pearson correlation was calculated for RPKM values of RNA-sequencing and ATACSeq.

**Supplementary Figure 6.**
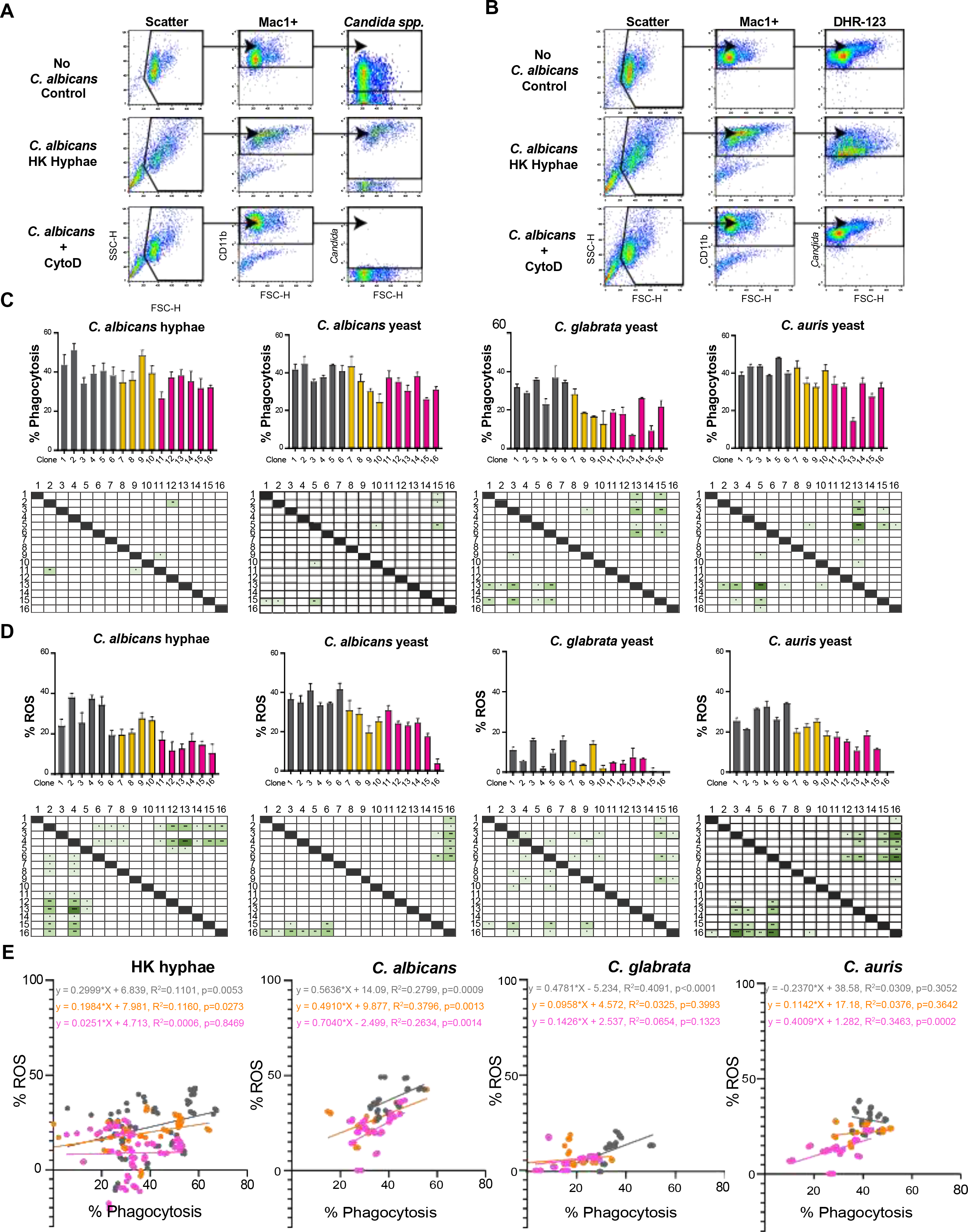
Functional characterization of clones through cytometry-based phagocytosis and ROS production assays. Neutrophils were co-incubated with a MOI 5 heat killed *C. albicans* hyphae for 2 hrs or MOI 3 heat killed *C. albicans* yeast, *C. glabrata*, or *C. auris* for 5 hrs at 37°C with CO2. **(a)** Representative gating scheme of phagocytosed *C. albicans* hyphae and **(b)** gating scheme of ROS production as measured by dihydrorhodamine-123 (DHR-123). **(c)** Percent phagocytosis for each clone calculated by subtracting the average cytochalasin D (CytoD) from each clone. **(d)** Percent ROS production for each clone. For **(c-d)**, mean with SEM plotted for each clone across 4 experiments for *C. albicans* hyphae and 2 experiments for *C. albicans* yeast, *C. glabrata*, and *C. auris* and samples were run as triplicate wells for each experiment. (e) Correlation tests for percent phagocytosis and percent ROS scores by functional cluster group. Stats: Kruskall-Wallis with Dunn’s post-test, * p ≤ 0.05, ** p ≤ 0.01, *** p ≤ 0.001, and **** p ≤ 0.001.

**Supplementary Figure 7.**
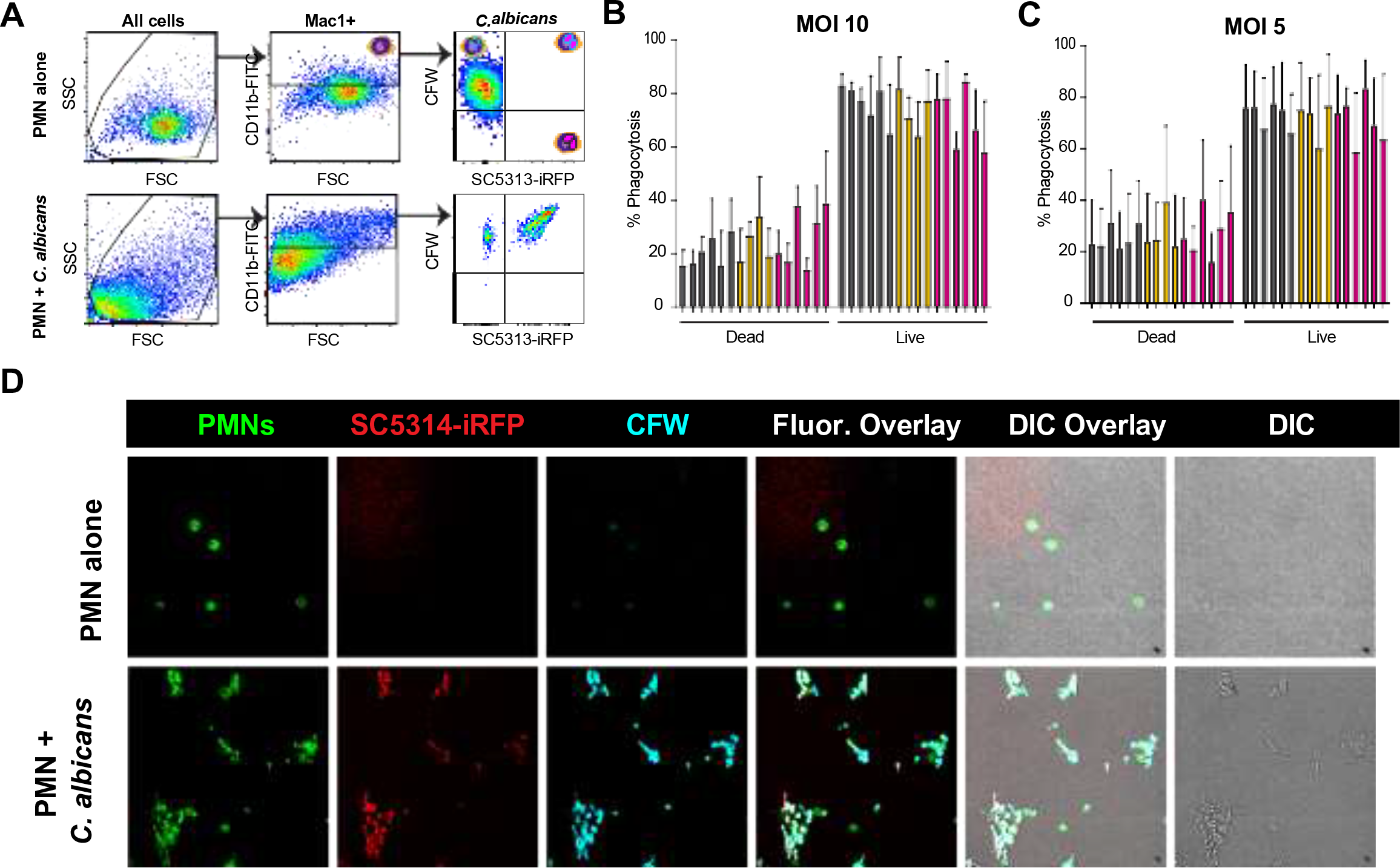
Functional characterization of clones through cytometry-based killing of wild type *C. albicans*. Neutrophils were co-incubated with a MOI 10 or MOI 5 of live far red fluorescent protein expressing wild type *C. albicans* (SC5314-iRFP) labelled with calcofluor white (CFW) for 2 hrs at 37°C. **(a)** Representative gating scheme of the flow-based fungal killing assay with anticipated live/dead outcomes for *C. albicans*. **(b)** Percent phagocytosed SC5314-iRPF cells at MOI 10 or **(c)** MOI 5 with live and dead yeast scores. Mean with SD plotted for each clone across 2 experiments and samples were run as one technical replicate. Stats: Kruskall-Wallis with Dunn’s post-test, all comparisons n.s. **(d)** Example fluorescence microscopy of each condition taken with confocal microscopy following co-incubation of samples.

**Supplementary Figure 8.**
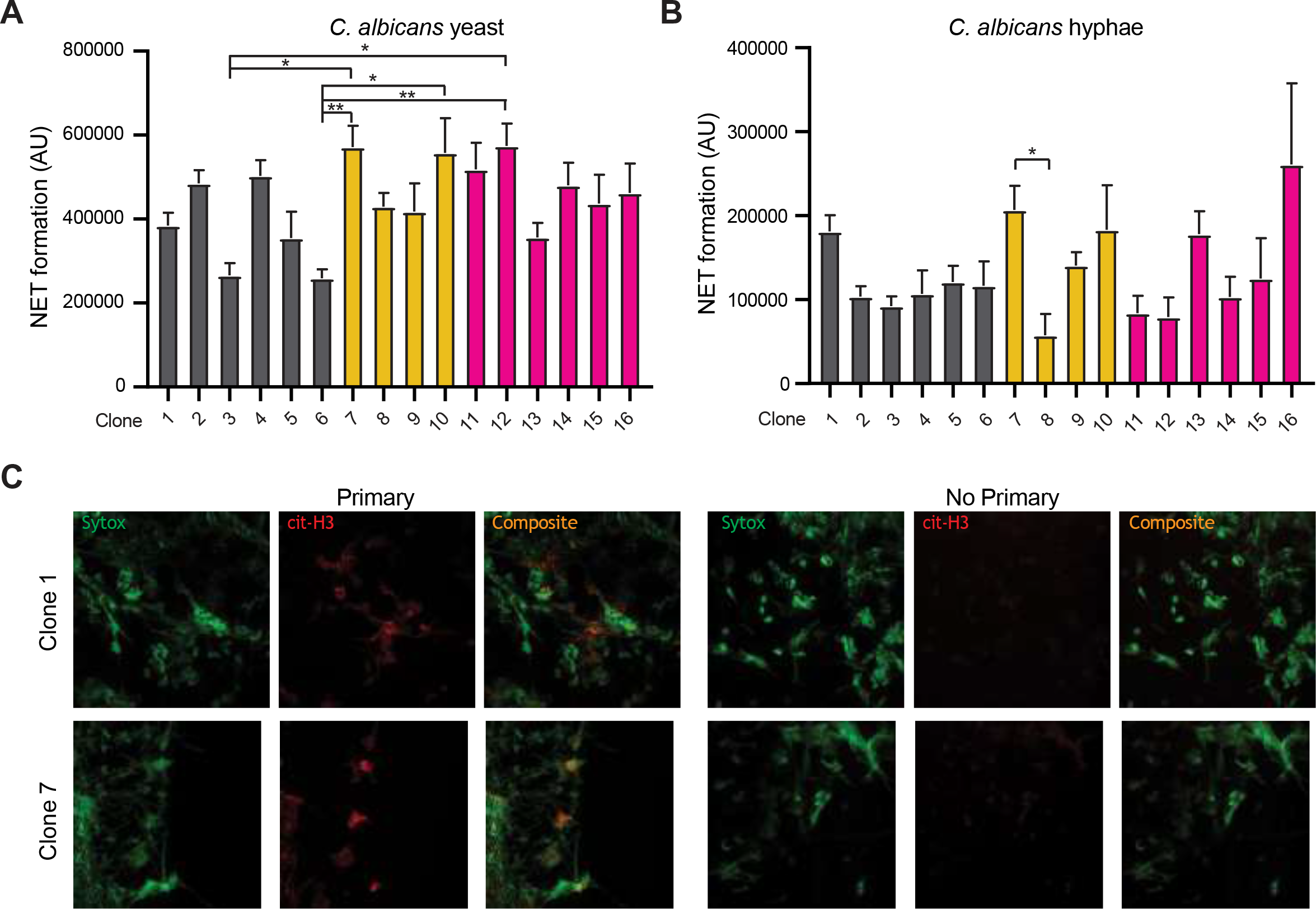
Formation of NETs between clonal neutrophils. Clones were coincubated for 5 hrs with Sytox green (final concentration 1 μM) and **(a)** *C. albicans* yeast at MOI 10 or **(b)** hyphae at roughly 3 × 108 cells/ml. Sytox fluorescence corresponding with extracellular DNA was measured in arbitrary units (AU). The average fluorescence from resting Neutrophils co-incubated in Sytox was subtracted from each *C. albicans* challenged well and A23187 was used as a positive control to normalize the data. Graphs are from three pooled experiments showing the mean and SEM. Samples were run as three technical replicates in each experiment. Stats: Kruskall-Wallis with Dunn’s post-test* p ≤ 0.05, ** p ≤ 0.01, *** p ≤ 0.001, and **** p ≤ 0.001. **(c)** Confocal microscopy was used to identify NET formation through counter immunostaining with anti-citrullinated-histone-specific antibody Cit-H3. Images are shown for two representative clones. Scale bar, 25 μm.

**Supplementary Figure 9.**
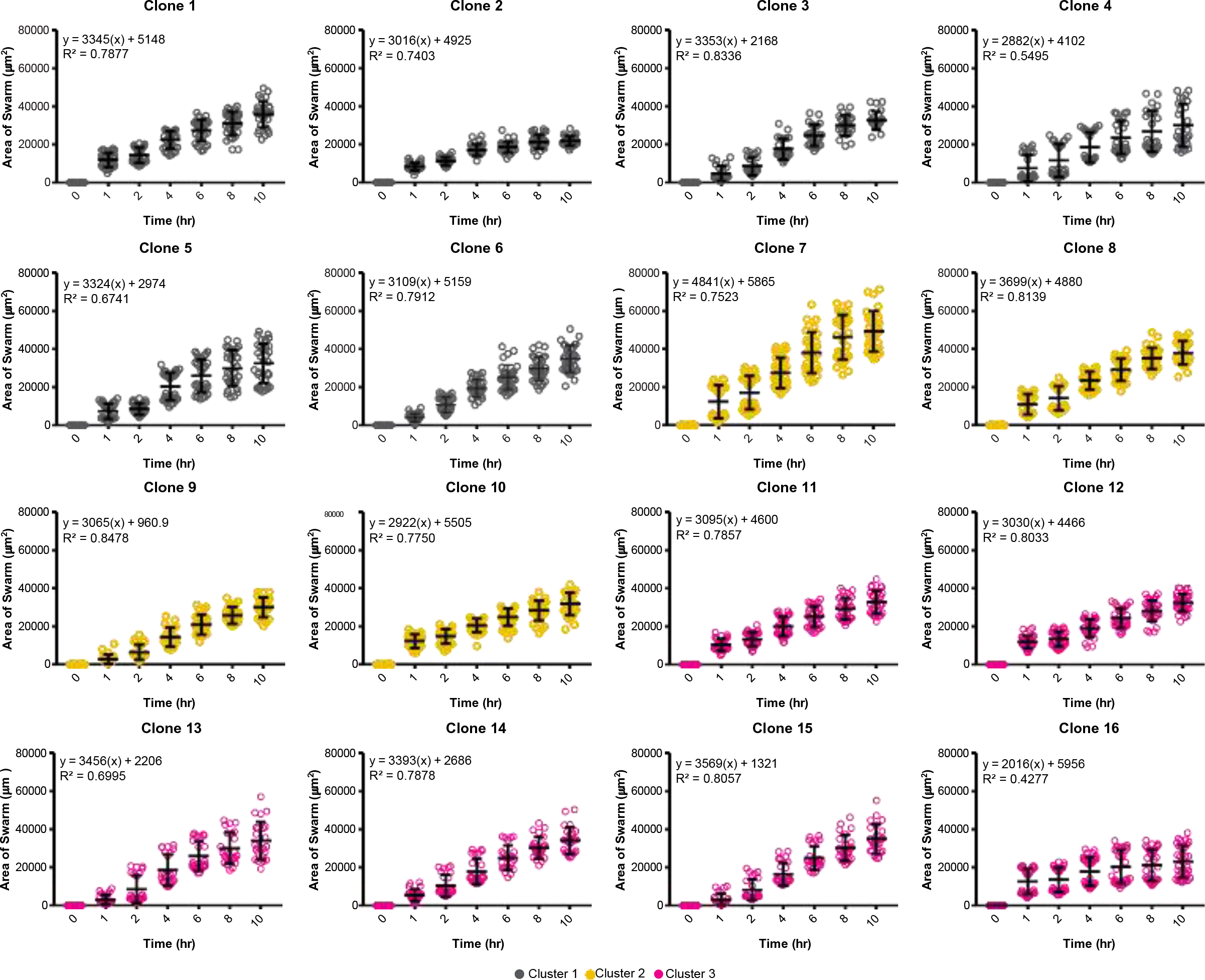
Swarming response to live *C. albicans* differs between clones over time. Neutrophil swarming response to SC5314-iRFP targets was quantified from microscopy over time. Swarming was measured by quantifying Hoescht-labelled neutrophils presented as the area of swarm over the course of 10 hrs. N = 2 independent experiments with n ≥ 23 swarms quantified per clone. A line of best fit was calculated for each clone and R2 values are provided.

**Supplementary Table 1.** P-values for the fraction of intersecting DEGs.

| Comparison | GMP vs GMP | PMN vs PMN | Common DEGs | p-value |
| --- | --- | --- | --- | --- |
| 1 vs 7 | 243 | 192 | 25 | 3.09E-25 |
| 1 vs 8 | 557 | 576 | 80 | 7.15E-53 |
| 1 vs 11 | 763 | 967 | 178 | 6.29E-118 |
| 1 vs 13 | 1148 | 1061 | 259 | 2.34E-160 |
| 1 vs 14 | 630 | 526 | 120 | 4.18E-100 |
| 1 vs 4 | 688 | 628 | 104 | 1.97E-67 |
| 1 vs 16 | 1012 | 873 | 240 | 6.98E-175 |
| 7 vs 8 | 677 | 452 | 105 | 8.86E-85 |
| 7 vs 11 | 769 | 611 | 161 | 5.74E-132 |
| 7 vs 13 | 1068 | 1061 | 252 | 7.96E-161 |
| 7 vs 14 | 558 | 459 | 116 | 9.04E-109 |
| 7 vs 4 | 574 | 342 | 85 | 2.29E-77 |
| 7 vs 16 | 926 | 1173 | 236 | 1.10E-147 |
| 8 vs 11 | 649 | 629 | 137 | 5.10E-110 |
| 8 vs 13 | 1066 | 995 | 238 | 2.64E-152 |
| 8 vs 14 | 1003 | 566 | 197 | 1.40E-165 |
| 8 vs 4 | 1050 | 650 | 183 | 1.25E-130 |
| 8 vs 16 | 1249 | 1310 | 284 | 3.12E-151 |
| 11 vs 13 | 809 | 1079 | 182 | 5.30E-109 |
| 11 vs 14 | 890 | 827 | 198 | 1.33E-141 |
| 11 vs 4 | 844 | 476 | 124 | 5.78E-95 |
| 11 vs 16 | 1022 | 1669 | 271 | 2.77E-134 |
| 13 vs 14 | 1039 | 936 | 272 | 1.05E-203 |
| 13 vs 4 | 938 | 1120 | 222 | 1.43E-135 |
| 13 vs 16 | 941 | 1019 | 259 | 5.72E-189 |
| 14 vs 4 | 565 | 524 | 97 | 6.02E-76 |
| 14 vs 16 | 772 | 803 | 200 | 4.57E-160 |
| 4 vs 16 | 688 | 1491 | 179 | 2.49E-94 |

**Supplementary Table 2.**
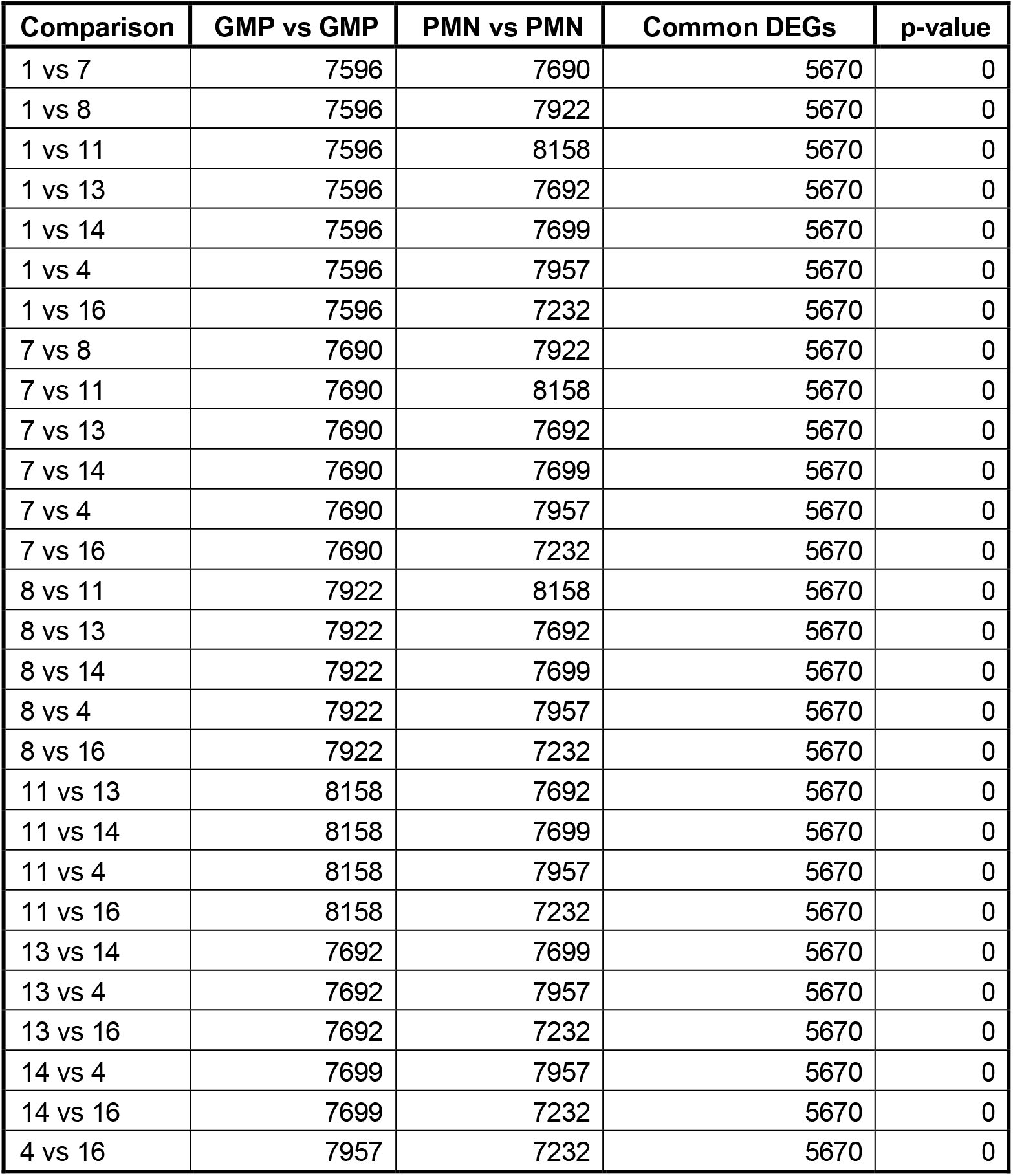
P-values for the fraction for all common DEGs.

| Comparison | GMP vs GMP | PMN vs PMN | Common DEGs | p-value |
| --- | --- | --- | --- | --- |
| 1 vs 7 | 7596 | 7690 | 5670 | 0 |
| 1 vs 8 | 7596 | 7922 | 5670 | 0 |
| 1 vs 11 | 7596 | 8158 | 5670 | 0 |
| 1 vs 13 | 7596 | 7692 | 5670 | 0 |
| 1 vs 14 | 7596 | 7699 | 5670 | 0 |
| 1 vs 4 | 7596 | 7957 | 5670 | 0 |
| 1 vs 16 | 7596 | 7232 | 5670 | 0 |
| 7 vs 8 | 7690 | 7922 | 5670 | 0 |
| 7 vs 11 | 7690 | 8158 | 5670 | 0 |
| 7 vs 13 | 7690 | 7692 | 5670 | 0 |
| 7 vs 14 | 7690 | 7699 | 5670 | 0 |
| 7 vs 4 | 7690 | 7957 | 5670 | 0 |
| 7 vs 16 | 7690 | 7232 | 5670 | 0 |
| 8 vs 11 | 7922 | 8158 | 5670 | 0 |
| 8 vs 13 | 7922 | 7692 | 5670 | 0 |
| 8 vs 14 | 7922 | 7699 | 5670 | 0 |
| 8 vs 4 | 7922 | 7957 | 5670 | 0 |
| 8 vs 16 | 7922 | 7232 | 5670 | 0 |
| 11 vs 13 | 8158 | 7692 | 5670 | 0 |
| 11 vs 14 | 8158 | 7699 | 5670 | 0 |
| 11 vs 4 | 8158 | 7957 | 5670 | 0 |
| 11 vs 16 | 8158 | 7232 | 5670 | 0 |
| 13 vs 14 | 7692 | 7699 | 5670 | 0 |
| 13 vs 4 | 7692 | 7957 | 5670 | 0 |
| 13 vs 16 | 7692 | 7232 | 5670 | 0 |
| 14 vs 4 | 7699 | 7957 | 5670 | 0 |
| 14 vs 16 | 7699 | 7232 | 5670 | 0 |
| 4 vs 16 | 7957 | 7232 | 5670 | 0 |

## REFERENCES

Accarias, S., T. Sanchez, A. Labrousse, M. Ben-Neji, A. Boyance, R. Poincloux, I. Maridonneau-Parini, and V.L. Cabec. 2020. Genetic engineering of hoxb8 immortalized hematopoietic progenitors: a potent tool to study macrophage tissue migration. Journal of Cell Science. jcs.236703. doi:10.1242/jcs.236703.

Anders, S., P.T. Pyl, and W. Huber. 2015. HTSeq—a Python framework to work with high-throughput sequencing data. Bioinformatics. 31:166–169. doi:10.1093/bioinformatics/btu638.

Bailey, T.L., J. Johnson, C.E. Grant, and W.S. Noble. 2015. The MEME Suite. Nucleic Acids Res. 43:W39–W49. doi:10.1093/nar/gkv416.

Bashant, K.R., A.M. Aponte, D. Randazzo, P.R. Sangsari, A.J. Wood, J.A. Bibby, E.E. West, A. Vassallo, Z.G. Manna, M.P. Playford, N. Jordan, S. Hasni, M. Gucek, C. Kemper, A.C. Morris, N.Y. Morgan, N. Toepfner, J. Guck, N.N. Mehta, E.R. Chilvers, C. Summers, and M.J. Kaplan. 2021. Proteomic, biomechanical and functional analyses define neutrophil heterogeneity in systemic lupus erythematosus. Ann Rheum Dis. 80:209–218. doi:10.1136/annrheumdis-2020-218338.

Basu, S., G. Hodgson, M. Katz, and A.R. Dunn. 2002. Evaluation of role of G-CSF in the production, survival, and release of neutrophils from bone marrow into circulation. Blood. 100:854–861. doi:10.1182/blood.v100.3.854.

Buenrostro, J.D., B. Wu, H.Y. Chang, and W.J. Greenleaf. 2015. ATAC-seq:A Method for Assaying Chromatin Accessibility Genome-Wide. Curr Protoc Mol Biology. 109:21.29.1–21.29.9. doi:10.1002/0471142727.mb2129s109.

Cabron, A.-S., K.E. Azzouzi, M. Boss, P. Arnold, J. Schwarz, M. Rosas, J.P. Dobert, E. Pavlenko, N. Schumacher, T. Renné, P.R. Taylor, S. Linder, S. Rose-John, and F. Zunke. 2018. Structural and Functional Analyses of the Shedding Protease ADAM17 in HoxB8-Immortalized Macrophages and Dendritic-like Cells. The Journal of Immunology. 201:ji1701556-3118. doi:10.4049/jimmunol.1701556.

Challen, G.A., N. Boles, K.K. Lin, and M.A. Goodell. 2009. Mouse hematopoietic stem cell identification and analysis. Cytom Part A. 75A:14–24. doi:10.1002/cyto.a.20674.

Chen, E.Y., C.M. Tan, Y. Kou, Q. Duan, Z. Wang, G.V. Meirelles, N.R. Clark, and A. Ma’ayan. 2013. Enrichr:interactive and collaborative HTML5 gene list enrichment analysis tool. Bmc Bioinformatics. 14:128–128. doi:10.1186/1471-2105-14-128.

Chu, J.Y., B. McCormick, G. Mazelyte, M. Michael, and S. Vermeren. 2018. HoxB8 neutrophils replicate Fcγ receptor and integrin-induced neutrophil signaling and functions. Journal of leukocyte biology. doi:10.1002/jlb.1ab0618-232r.

Deerhake, M.E., E.Y. Reyes, S. Xu-Vanpala, and M.L. Shinohara. 2021. Single-Cell Transcriptional Heterogeneity of Neutrophils During Acute Pulmonary Cryptococcus neoformans Infection. Front Immunol. 12:670574. doi:10.3389/fimmu.2021.670574.

Dobin, A., C.A. Davis, F. Schlesinger, J. Drenkow, C. Zaleski, S. Jha, P. Batut, M. Chaisson, and T.R. Gingeras. 2013. STAR:ultrafast universal RNA-seq aligner. Bioinformatics. 29:15–21. doi:10.1093/bioinformatics/bts635.

Filep, J.G., and A. Ariel. 2020. Neutrophil heterogeneity and fate in inflamed tissues:implications for the resolution of inflammation. Am J Physiol-cell Ph. 319:C510–C532. doi:10.1152/ajpcell.00181.2020.

Fites, J.S., M. Gui, J.F. Kernien, P. Negoro, Z. Dagher, D.B. Sykes, J.E. Nett, M.K. Mansour, and B.S. Klein. 2018. An unappreciated role for neutrophil-DC hybrids in immunity to invasive fungal infections. Plos Pathog. 14:e1007073. doi:10.1371/journal.ppat.1007073.

Friedman, A.D. 2007. Transcriptional control of granulocyte and monocyte development. Oncogene. 26:6816–6828. doi:10.1038/sj.onc.1210764.

Gazendam, R.P., J.L. van Hamme, A.T.J. Tool, M. van Houdt, P.J.J.H. Verkuijlen, M. Herbst, J.G. Liese, F.L. van de Veerdonk, D. Roos, T.K. van den Berg, and T.W. Kuijpers. 2014. Two independent killing mechanisms of Candida albicans by human neutrophils:evidence from innate immunity defects. Blood. 124:590–597. doi:10.1182/blood-2014-01-551473.

Grieshaber-Bouyer, R., and P.A. Nigrovic. 2019. Neutrophil Heterogeneity as Therapeutic Opportunity in Immune-Mediated Disease. Front Immunol. 10:346. doi:10.3389/fimmu.2019.00346.

Guilliams, M., A. Mildner, and S. Yona. 2018. Developmental and Functional Heterogeneity of Monocytes. Immunity. 49:595–613. doi:10.1016/j.immuni.2018.10.005.

Gurzeler, U., T. Rabachini, C.A. Dahinden, M. Salmanidis, G. Brumatti, P.G. Ekert, N. Echeverry, D. Bachmann, H.U. Simon, and T. Kaufmann. 2013. In vitro differentiation of near-unlimited numbers of functional mouse basophils using conditional Hoxb8. Allergy. 68:604–613. doi:10.1111/all.12140.

Hellebrekers, P., F. Hietbrink, N. Vrisekoop, L.P.H. Leenen, and L. Koenderman. 2017. Neutrophil Functional Heterogeneity:Identification of Competitive Phagocytosis. Front Immunol. 8:1498. doi:10.3389/fimmu.2017.01498.

Hopke, A., N. Nicke, E.E. Hidu, G. Degani, L. Popolo, and R.T. Wheeler. 2016. Neutrophil Attack Triggers Extracellular Trap-Dependent Candida Cell Wall Remodeling and Altered Immune Recognition. Plos Pathog. 12:e1005644. doi:10.1371/journal.ppat.1005644.

Hopke, A., A. Scherer, S. Kreuzburg, M.S. Abers, C.S. Zerbe, M.C. Dinauer, M.K. Mansour, and D. Irimia. 2020. Neutrophil swarming delays the growth of clusters of pathogenic fungi. Nat Commun. 11:2031. doi:10.1038/s41467-020-15834-4.

Hopke, A., and R. Wheeler. 2017. In vitro Detection of Neutrophil Traps and Post-attack Cell Wall Changes in Candida Hyphae. Bio-protocol. 7. doi:10.21769/bioprotoc.2213.

Huang, D.W., B.T. Sherman, and R.A. Lempicki. 2009. Bioinformatics enrichment tools:paths toward the comprehensive functional analysis of large gene lists. Nucleic Acids Res. 37:1–13. doi:10.1093/nar/gkn923.

John, S., P.J. Sabo, R.E. Thurman, M.-H. Sung, S.C. Biddie, T.A. Johnson, G.L. Hager, and J.A. Stamatoyannopoulos. 2011. Chromatin accessibility pre-determines glucocorticoid receptor binding patterns. Nat Genet. 43:264–268. doi:10.1038/ng.759.

Johnson, K.D., D.J. Conn, E. Shishkova, K.R. Katsumura, P. Liu, S. Shen, E.A. Ranheim, S.G. Kraus, W. Wang, K.R. Calvo, A.P. Hsu, S.M. Holland, J.J. Coon, S. Keles, and E.H. Bresnick. 2020. Constructing and deconstructing GATA2-regulated cell fate programs to establish developmental trajectories. J Exp Med. 217:e20191526. doi:10.1084/jem.20191526.

Johnson, W.E., C. Li, and A. Rabinovic. 2007. Adjusting batch effects in microarray expression data using empirical Bayes methods. Biostatistics. 8:118–127. doi:10.1093/biostatistics/kxj037.

Katzmarski, N., J. Domínguez-Andrés, B. Cirovic, G. Renieris, E. Ciarlo, D.L. Roy, K. Lepikhov, K. Kattler, G. Gasparoni, K. Händler, H. Theis, M. Beyer, J.W.M. van der Meer, L.A.B. Joosten, J. Walter, J.L. Schultze, T. Roger, E.J. Giamarellos-Bourboulis, A. Schlitzer, and M.G. Netea. 2021. Transmission of trained immunity and heterologous resistance to infections across generations. Nat Immunol. 1–9. doi:10.1038/s41590-021-01052-7.

Kenny, E.F., A. Herzig, R. Krüger, A. Muth, S. Mondal, P.R. Thompson, V. Brinkmann, H. von Bernuth, and A. Zychlinsky. 2017. Diverse stimuli engage different neutrophil extracellular trap pathways. Elife. 6:e24437. doi:10.7554/elife.24437.

Kuleshov, M.V., M.R. Jones, A.D. Rouillard, N.F. Fernandez, Q. Duan, Z. Wang, S. Koplev, S.L. Jenkins, K.M. Jagodnik, A. Lachmann, M.G. McDermott, C.D. Monteiro, G.W. Gundersen, and A. Ma’ayan. 2016. Enrichr:a comprehensive gene set enrichment analysis web server 2016 update. Nucleic Acids Res. 44:W90–W97. doi:10.1093/nar/gkw377.

Kwok, I., E. Becht, Y. Xia, M. Ng, Y.C. Teh, L. Tan, M. Evrard, J.L.Y. Li, H.T.N. Tran, Y. Tan, D. Liu, A. Mishra, K.H. Liong, K. Leong, Y. Zhang, A. Olsson, C.K. Mantri, P. Shyamsunder, Z. Liu, C. Piot, C.-A. Dutertre, H. Cheng, S. Bari, N. Ang, S.K. Biswas, H.P. Koeffler, H.L. Tey Larbi, I.-H. Su, B. Lee, A.St. John, J.K.Y. Chan, W.Y.K. Hwang, J. Chen, N. Salomonis, S.Z. Chong, H.L. Grimes, B. Liu, A. Hidalgo, E.W. Newell, T. Cheng, F. Ginhoux, and L.G. Ng. 2020. Combinatorial Single-Cell Analyses of Granulocyte-Monocyte Progenitor Heterogeneity Reveals an Early Uni-potent Neutrophil Progenitor. Immunity. 53:303–318.e5. doi:10.1016/j.immuni.2020.06.005.

Leek, J.T., W.E. Johnson, H.S. Parker, A.E. Jaffe, and J.D. Storey. 2012. The sva package for removing batch effects and other unwanted variation in high-throughput experiments. Bioinformatics. 28:882–883. doi:10.1093/bioinformatics/bts034.

Li, G., W. Hao, and W. Hu. 2020. Transcription factor PU.1 and immune cell differentiation (Review). Int J Mol Med. 46:1943–1950. doi:10.3892/ijmm.2020.4763.

Li, H., and R. Durbin. 2009. Fast and accurate short read alignment with Burrows–Wheeler transform. Bioinformatics. 25:1754–1760. doi:10.1093/bioinformatics/btp324.

Ma, G., D. Gezer, O. Herrmann, K. Feldberg, M. Schemionek, M. Jawhar, A. Reiter, T.H. Brümmendorf, S. Koschmieder, and N. Chatain. 2020. LCP1 triggers mTORC2/AKT activity and is pharmacologically targeted by enzastaurin in hypereosinophilia. Molecular carcinogenesis. 59:87–103. doi:10.1002/mc.23131.

Maskarinec, S.A., M. McKelvy, K. Boyle, H. Hotchkiss, M.E. Duarte, B. Addison, N. Amato, S. Khandelwal, G.M. Arepally, and G.M. Lee. 2022. Neutrophil functional heterogeneity is a fixed phenotype and is associated with distinct gene expression profiles. J Leukocyte Biol. doi:10.1002/jlb.4a0322-164r.

Mercier, F.E., D.B. Sykes, and D.T. Scadden. 2016. Single Targeted Exon Mutation Creates a True Congenic Mouse for Competitive Hematopoietic Stem Cell Transplantation:The C57BL/6-CD45.1STEM Mouse. Stem Cell Rep. 6:985–992. doi:10.1016/j.stemcr.2016.04.010.

Mistry, P., S. Nakabo, L. O’Neil, R.R. Goel, K. Jiang, C. Carmona-Rivera, S. Gupta, D.W. Chan, P.M. Carlucci, X. Wang, F. Naz, Z. Manna, A. Dey, N.N. Mehta, S. Hasni, S. Dell’Orso, G. Gutierrez-Cruz, H.-W. Sun, and M.J. Kaplan. 2019. Transcriptomic, epigenetic, and functional analyses implicate neutrophil diversity in the pathogenesis of systemic lupus erythematosus. Proc National Acad Sci. 116:25222–25228. doi:10.1073/pnas.1908576116.

Negoro, P.E., S. Xu, Z. Dagher, A. Hopke, J.L. Reedy, M.B. Feldman, N.S. Khan, A.L. Viens, N.J. Alexander, N.J. Atallah, A.K. Scherer, R.A. Dutko, J. Jeffery, J.F. Kernien, J.S. Fites, J.E. Nett, B.S. Klein, J.M. Vyas, D. Irimia, D.B. Sykes, and M.K. Mansour. 2020. Spleen Tyrosine Kinase Is a Critical Regulator of Neutrophil Responses to Candida Species. Mbio. 11:e02043-19. doi:10.1128/mbio.02043-19.

Orosz, A., B. Walzog, and A. Mócsai. 2020. In Vivo Functions of Mouse Neutrophils Derived from HoxB8-Transduced Conditionally Immortalized Myeloid Progenitors. J Immunol. ji2000807. doi:10.4049/jimmunol.2000807.

Pillay, J., I. den Braber, N. Vrisekoop, L.M. Kwast, R.J. de Boer, J.A.M. Borghans, K. Tesselaar, and L. Koenderman. 2010. In vivo labeling with 2H2O reveals a human neutrophil lifespan of 5.4 days. Blood. 116:625–627. doi:10.1182/blood-2010-01-259028.

Poplimont, H., A. Georgantzoglou, M. Boulch, H.A. Walker, C. Coombs, F. Papaleonidopoulou, and M. Sarris. 2020. Neutrophil Swarming in Damaged Tissue Is Orchestrated by Connexins and Cooperative Calcium Alarm Signals. Curr Biol. 30:2761–2776.e7. doi:10.1016/j.cub.2020.05.030.

Qi, X., Y. Yu, R. Sun, J. Huang, L. Liu, Y. Yang, T. Rui, and B. Sun. 2021. Identification and characterization of neutrophil heterogeneity in sepsis. Crit Care. 25:50. doi:10.1186/s13054-021-03481-0.

Ricotta, E.E., Y.L. Lai, A. Babiker, J.R. Strich, S.S. Kadri, M.S. Lionakis, D.R. Prevots, and J. Adjemian. 2020. Invasive candidiasis species distribution and trends, United States, 2009-2017. J Infect Dis. 223:jiaa502-. doi:10.1093/infdis/jiaa502.

Robinson, M.D., D.J. McCarthy, and G.K. Smyth. 2010. edgeR:a Bioconductor package for differential expression analysis of digital gene expression data. Bioinformatics. 26:139–140. doi:10.1093/bioinformatics/btp616.

Rodrigues, N.P., A.S. Boyd, C. Fugazza, G.E. May, Y. Guo, A.J. Tipping, D.T. Scadden, P. Vyas, and T. Enver. 2008. GATA-2 regulates granulocyte-macrophage progenitor cell function. Blood. 112:4862–4873. doi:10.1182/blood-2008-01-136564.

Ross-Innes, C.S., R. Stark, A.E. Teschendorff, K.A. Holmes, H.R. Ali, M.J. Dunning, G.D. Brown, O. Gojis, I.O. Ellis, A.R. Green, S. Ali, S.-F. Chin, C. Palmieri, C. Caldas, and J.S. Carroll. 2012. Differential oestrogen receptor binding is associated with clinical outcome in breast cancer. Nature. 481:389–393. doi:10.1038/nature10730.

Saul, S., C. Castelbou, C. Fickentscher, and N. Demaurex. 2019. Signaling and functional competency of neutrophils derived from bone-marrow cells expressing the ER-HOXB8 oncoprotein. Journal of leukocyte biology. doi:10.1002/jlb.2a0818-314r.

Scherer, A.K., A. Hopke, D.B. Sykes, D. Irimia, and M.K. Mansour. 2021. Host defense against fungal pathogens:Adaptable neutrophil responses and the promise of therapeutic opportunities? Plos Pathog. 17:e1009691. doi:10.1371/journal.ppat.1009691.

Shankar, M., T.L. Lo, and A. Traven. 2020. Natural Variation in Clinical Isolates of Candida albicans Modulates Neutrophil Responses. Msphere. 5:e00501–20. doi:10.1128/msphere.00501-20.

Sollberger, G., D.O. Tilley, and A. Zychlinsky. 2018. Neutrophil Extracellular Traps:The Biology of Chromatin Externalization. Dev Cell. 44:542–553. doi:10.1016/j.devcel.2018.01.019.

Stark, R., and G. Brown. 8AD. DiffBind:Differential binding analysis of ChIPSeq peak data.

Sykes, D.B., and M.P. Kamps. 2001. Estrogen-dependent E2a/Pbx1 myeloid cell lines exhibit conditional differentiation that can be arrested by other leukemic oncoproteins. Blood. 98:2308–2318. doi:10.1182/blood.v98.8.2308.

Sykes, D.B., Y.S. Kfoury, F.E. Mercier, M.J. Wawer, J.M. Law, M.K. Haynes, T.A. Lewis, A. Schajnovitz, E. Jain, D. Lee, H. Meyer, K.A. Pierce, N.J. Tolliday, A. Waller, S.J. Ferrara, A.L. Eheim, D. Stoeckigt, K.L. Maxcy, J.M. Cobert, J. Bachand, B.A. Szekely, S. Mukherjee, L.A. Sklar, J.D. Kotz, C.B. Clish, R.I. Sadreyev, P.A. Clemons, A. Janzer, S.L. Schreiber, and D.T. Scadden. 2016. Inhibition of Dihydroorotate Dehydrogenase Overcomes Differentiation Blockade in Acute Myeloid Leukemia. Cell. 167:171–186.e15. doi:10.1016/j.cell.2016.08.057.

Tak, T., K. Tesselaar, J. Pillay, J.A.M. Borghans, and L. Koenderman. 2013. Whatˈs your age again? Determination of human neutrophil half-lives revisited. J Leukocyte Biol. 94:595–601. doi:10.1189/jlb.1112571.

Urban, C.F., and E. Backman. 2020. Eradicating, retaining, balancing, swarming, shuttling and dumping:a myriad of tasks for neutrophils during fungal infection. Curr Opin Microbiol. 58:106–115. doi:10.1016/j.mib.2020.09.011.

Villani, A.-C., R. Satija, G. Reynolds, S. Sarkizova, K. Shekhar, J. Fletcher, M. Griesbeck, A. Butler, S. Zheng, S. Lazo, L. Jardine, D. Dixon, E. Stephenson, E. Nilsson, I. Grundberg, D. McDonald, A. Filby, W. Li, P.L.D. Jager, O. Rozenblatt-Rosen, A.A. Lane, M. Haniffa, A. Regev, and N. Hacohen. 2017. Single-cell RNA-seq reveals new types of human blood dendritic cells, monocytes, and progenitors. Science. 356:eaah4573. doi:10.1126/science.aah4573.

Vizcaíno, C., S. Mansilla, and J. Portugal. 2015. Sp1 transcription factor:A long-standing target in cancer chemotherapy. Pharmacol Therapeut. 152:111–124. doi:10.1016/j.pharmthera.2015.05.008.

Wang, G.G., K.R. Calvo, M.P. Pasillas, D.B. Sykes, H. Häcker, and M.P. Kamps. 2006. Quantitative production of macrophages or neutrophils ex vivo using conditional Hoxb8. Nat Methods. 3:287–293. doi:10.1038/nmeth865.

Wolf, A.A., A. Yáñez, P.K. Barman, and H.S. Goodridge. 2019. The Ontogeny of Monocyte Subsets. Front Immunol. 10:1642. doi:10.3389/fimmu.2019.01642.

Xie, X., Q. Shi, P. Wu, X. Zhang, H. Kambara, J. Su, H. Yu, S.-Y. Park, R. Guo, Q. Ren, S. Zhang, Y. Xu, L.E. Silberstein, T. Cheng, F. Ma, C. Li, and H.R. Luo. 2020. Single-cell transcriptome profiling reveals neutrophil heterogeneity in homeostasis and infection. Nat Immunol. 21:1119–1133. doi:10.1038/s41590-020-0736-z.

Xie, Z., A. Bailey, M.V. Kuleshov, D.J.B. Clarke, J.E. Evangelista, S.L. Jenkins, A. Lachmann, M.L. Wojciechowicz, E. Kropiwnicki, K.M. Jagodnik, M. Jeon, and A. Ma’ayan. 2021. Gene Set Knowledge Discovery with Enrichr. Curr Protoc. 1:e90. doi:10.1002/cpz1.90.

Yáñez, A., M.Y. Ng, N. Hassanzadeh-Kiabi, and H.S. Goodridge. 2015. IRF8 acts in lineage-committed rather than oligopotent progenitors to control neutrophil vs monocyte production. Blood. 125:1452–1459. doi:10.1182/blood-2014-09-600833.

